# Flexible decisions arise from resource-rational memory sampling

**DOI:** 10.64898/2026.06.15.732446

**Authors:** Jonathan Nicholas, Sixing Chen, Marcelo G. Mattar

## Abstract

Flexible decision making depends on retrieving and recombining memories. Yet because this process unfolds covertly, its governing principles remain unknown. Here we use gaze reinstatement to uncover the hidden dynamics and computational logic of memory retrieval during flexible behavior. As people deliberated on a blank screen, they directed their gaze toward the encoding locations of decision-relevant experiences, and these fixations shaped their evolving choice. A task-optimized recurrent neural network captured both their behavior and gaze patterns by learning to balance retrieval costs against expected gains in decision quality. These results demonstrate that flexible decisions emerge from a resource-rational process in which memories are sampled to construct decision variables on the fly.

---

To make successful choices we must first know what our options are worth. But what matters often changes from one decision to the next, requiring us to construct value anew each time. Our episodic memory system enables this type of flexible valuation by allowing us to retrieve detailed experiences and recombine them as needed (1, 2). This capacity has been implicated in decision contexts ranging from simple choice to multistep planning and inference (3–10). Yet despite considerable progress in understanding *why* episodic memory supports adaptive behavior, far less is known about *how* it is recruited to guide choice. In particular, we lack a principled account of the algorithms that govern both which memories are retrieved and when they are accessed during deliberation.

Decisions between familiar options whose values are already known, by contrast, are well described by models in which noisy evidence is sequentially sampled and accumulates in favor of one option over another (11–14). Recent work has shown that, because cognitive resources are limited, people deliberate in this setting by optimally balancing the cost of taking each new sample against its expected improvement to decision quality — a *resource-rational* strategy (15, 16). When faced with a novel choice, however, people must instead knit value together across separate past experiences (17–19). Deliberation in this case is thought to rely on an analogous process in which evidence is sampled from individual memories and accumulates toward a decision (20–22), possibly governed by this same cost-benefit logic. Consistent with this account, rodent hippocampal place cells represent candidate options during flexible decision making, and do so more often when choices are harder (23–25).

Direct evidence for such sampling in humans is lacking because the fine-grained dynamics of episodic retrieval are difficult to measure. Uncovering how episodic memory guides choice therefore requires access to this latent process as it unfolds throughout deliberation. Eye movements offer a potential window: even in the absence of visual input, gaze spontaneously returns to spatial locations previously associated with remembered information, a phenomenon called *looking at nothing* (26– 28). Fixations may therefore serve as a covert real-time measure of retrieval.

Here, we leverage this phenomenon to make the hidden process of memory-guided choice observable as it unfolds in real time. Using a task in which people must make decisions by flexibly combining information across multiple distinct memories, we track their gaze to identify which memories they retrieve and when. We then ask whether the way people sample their memories represents an efficient solution to the problem of deciding well when retrieval is costly. To answer this question, we optimize the performance of a recurrent neural network on the same task and find that it learns a resource-rational sampling strategy that reproduces both the memories people retrieve and the choices they make. Finally, we characterize the principles that govern how this sampler is initialized at the start of deliberation as well as the order in which samples are retrieved thereafter. Together, these results show that flexible decisions emerge from a resource-rational deliberation process in which value is actively built from episodic memory.

## Measuring episodic sampling during deliberation

Forty-three participants completed several rounds of a four-part task (**Fig. 1A**) designed to capture episodic retrieval during flexible decision making while their eye gaze was recorded. During the encoding phase, participants saw episodes consisting of an item and an associated reward, one at a time. Each episode was displayed in one of six possible locations around a circle, and items varied across several features (**Fig. 1B**). Following encoding, participants completed a 2-back working memory task to prevent active rehearsal. Participants next viewed a blank screen and made decisions based on the features of each item. On each choice, participants heard an offer consisting of a single feature (e.g., “animal”) and decided whether to take or leave it. Participants were informed that an offer’s value was the sum of rewards associated with offer-relevant items (e.g., the value of “animal” was the sum of all animal rewards), and that they should take positive offers and leave negative offers. Following the decision phase, we assessed participants’ memory by asking them to view the same blank screen and to first verbally recall the items they had seen, and then to recall the reward and location of each item.

**Figure 1:**
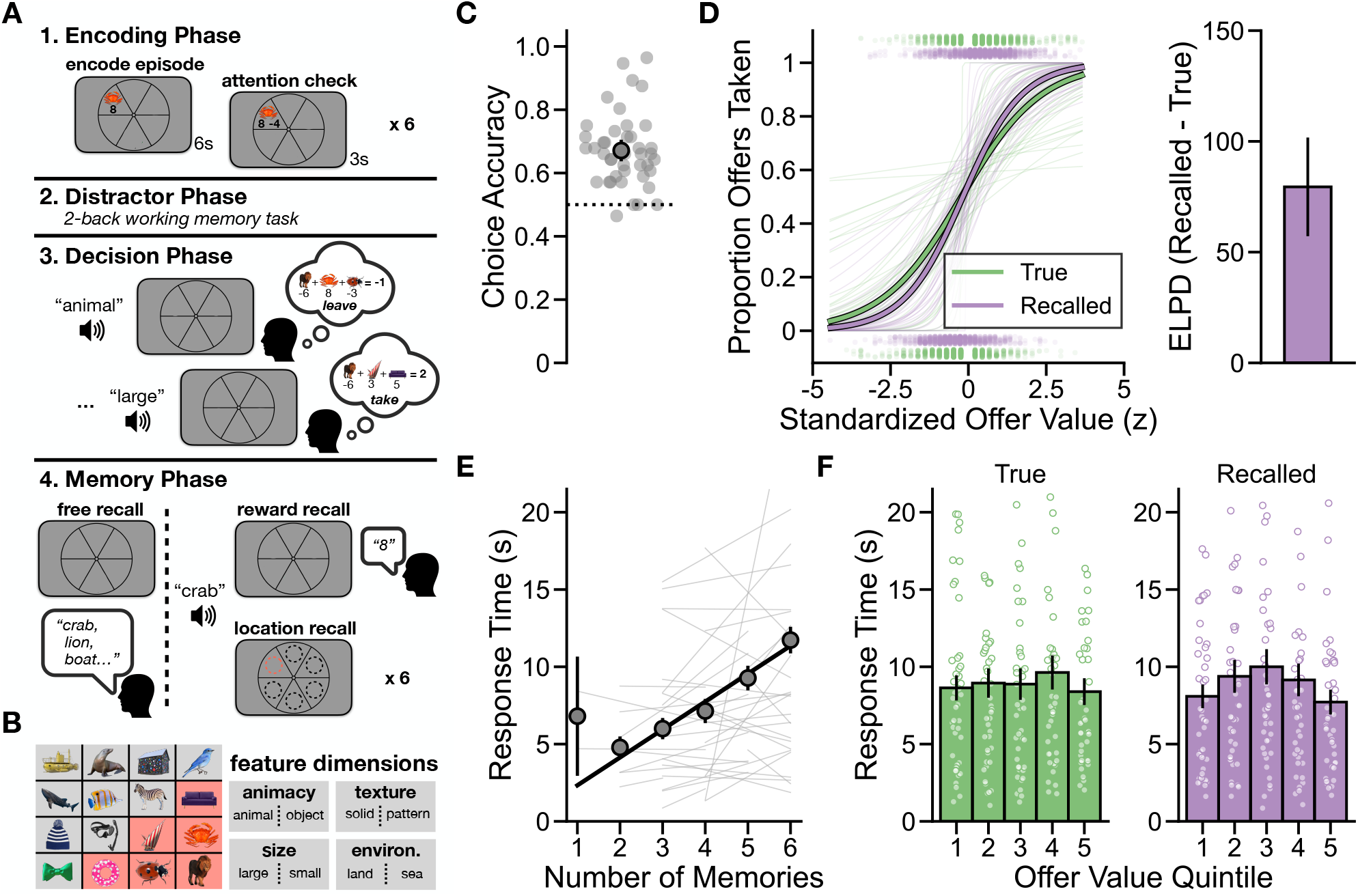
Experimental task and performance. **(A)** Each round consisted of four phases. Participants encoded six episodes at unique screen positions, each followed by an attention check in which they identified the item’s reward from two numbers: the true reward and a randomly selected foil. They then completed a 90-second 2-back working memory task, followed by the decision phase in which they viewed a blank screen and heard offers one at a time, with unlimited time to deliberate. Lastly, participants freely recalled the items from the round, and then heard each item one at a time and reported its reward and location **(B)** Left: the full set of images shown to participants. Six images were sampled to be shown in each round (an example six are highlighted in red). Right: The four binary features of each image. **(C)** Overall accuracy on the decision making phase. **(D)** Left: The proportion of offers that were taken as a function of summed true offer value and recalled offer value. Offer values are z-scored to facilitate comparison. Right: Predictive performance of the true and recalled offer value models. Expected log pointwise predictive density (ELPD) from ten-fold cross-validation. **(E)** Decision response times as a function of the number of memories accurately recalled on each round. **(F)** Decision response times as a function of true (left) and recalled (right) offer value, split into quintile bins. In all plots, individuals are shown as lighter points (averages) or lines (regression fits) with group-level averages (points) or regressions plotted in darker colors on top. Error bars represent standard error. Dotted horizontal lines indicate chance level.

Because each episode was relevant to some offers but not others, participants had to flexibly retrieve and recombine their memories to evaluate each offer. Further, because every episode was encoded at a distinct location, fixations to these locations provide a potential readout of which episodes were retrieved and when.

## Decisions show behavioral signatures of episodic memory

We first examined whether participants made effective decisions and recalled the episodes shown in each round. Participants performed well on the decision task, tending to take positive and to leave negative offers (*M*_*accuracy*_ = 67.1 ± 1.7%, *β*_0_ = 0.76, 95% *HDI* = [0.58, 0.94]; **Fig. 1C**). They also had robust episodic memory. Participants recalled an average of 77.1 ± 2.3% of the items that appeared in each round (**Fig. S1A**), and their memory for the rewards associated with each item was strongly related to the actual associated rewards (*β*_reward_ = 0.75, 95% *HDI* = [0.68, 0.82]; **Fig. S1B**). Finally, although the locations of each episode were incidental to the task, participants demonstrated highly accurate location memory (*M*_*accuracy*_ = 98.4 ± 0.5%, *β*_0_ = 0.82, 95% *HDI* = [0.81, 0.83]; **Fig. S1C**).

We next asked whether participants’ choices demonstrated signatures of episodic memory use during deliberation (1). If participants retrieve and sum over individual episodes at choice time, then their decision response times should scale with the number of episodes available for retrieval, and their choices should be better predicted by the summed value of remembered rewards (recalled offer value) than by the true offer value. We tested these predictions using free recall as an index of the number of episodes available during decision making, and by comparing the predictive power of recalled offer value versus true offer value.

Consistent with this account, participants took longer to make decisions on rounds in which they subsequently recalled more items (*β*_nMemories_ = 0.16, 95% *HDI* = [0.07, 0.26]; **Fig. 1E**) and their choices were better predicted out-of-sample by recalled offer value than by true offer value (*ELPD*_*recalled−true*_ = 79.46 ± 22.27; **Fig. 1D**; **Table S1**). In addition, participants took longer to make decisions with more ambiguous recalled offer values (i.e., closer to zero; 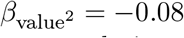, 95% HDI [−0.12, −0.04]; **Fig. 1F**), demonstrating a classic signature of evidence accumulation where harder choices require more samples to reach a decision threshold (11, 29). Together, these results indicate that participants relied on episodic retrieval to guide their decisions.

## Fixations capture sampling from episodic memory

Behavioral measures alone cannot reveal how retrieval unfolds throughout deliberation, so we next examined whether participants’ gaze tracked episodic retrieval. Before turning to decision making, where retrieval is hidden and its content is unknown, we first asked whether eye movements provide a reliable marker of episodic retrieval during the free recall task, where retrieval is instead explicit. We found that participants directed their gaze toward the encoding locations of items as they recalled them aloud (**Fig. 2A**), validating the use of eye movements as a covert marker of retrieval.

**Figure 2:**
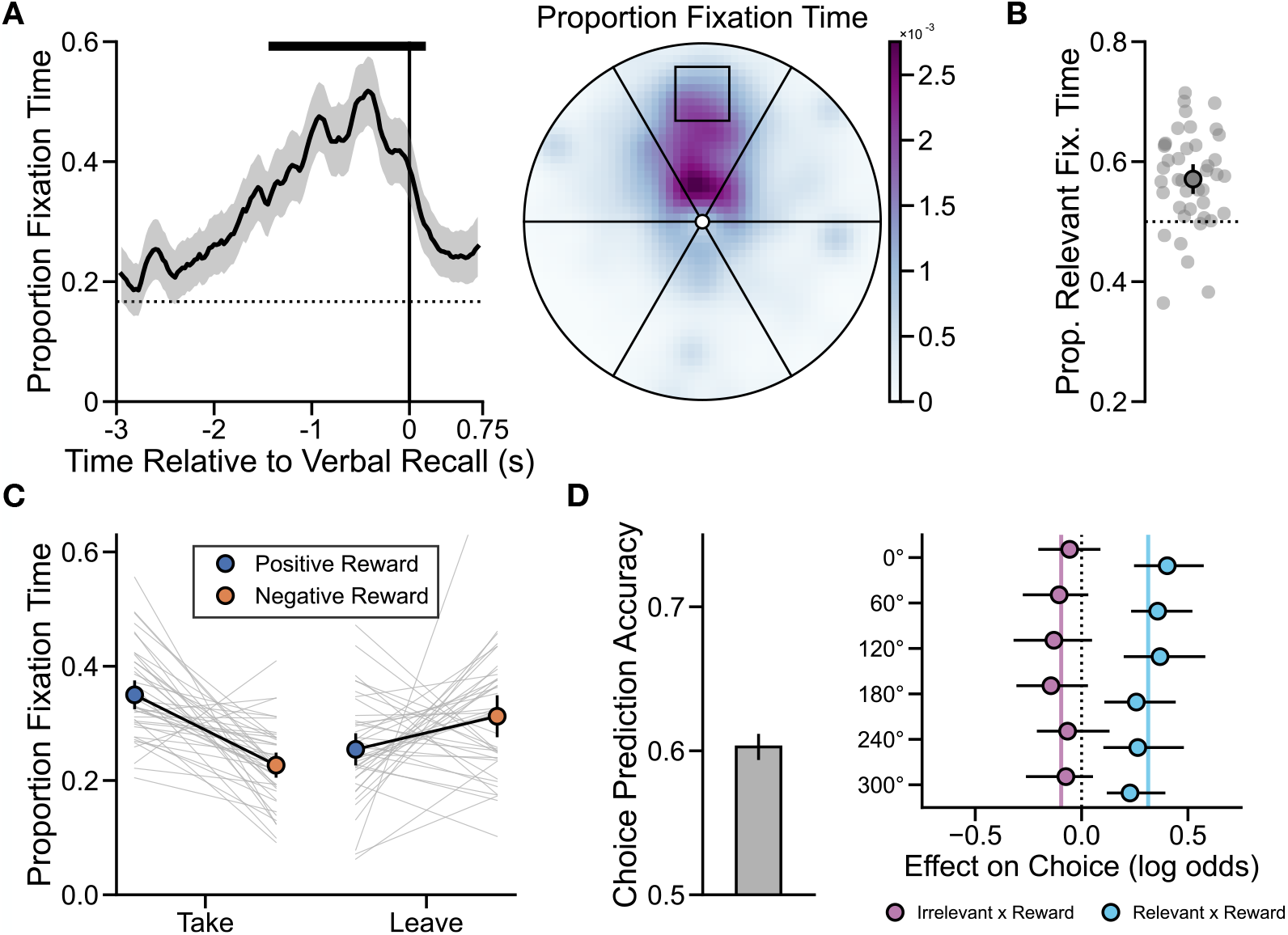
Gaze reflects episodic recall during decision making. **(A)** Left: Time course of proportion fixation time on the location of the item being verbally recalled during the free recall task. The line shows the group mean with standard error. The dotted line indicates chance and the vertical line indicates onset of verbal recall. The black bar above the trace marks the significant time points (cluster-corrected permutation test, *p* < 0.0001). Right: Spatial distribution of gaze to each item area of interest (“wedge” of the circle) during significant time points, displayed as a smoothed heatmap. Fixations for each recalled item were rotated as if the encoding location was at top-center (shown as a square) and proportions were calculated for each subject and then averaged. **(B)** Proportion of fixation time directed toward offer-relevant episode locations during decision making. Each point is one participant; group mean shown as the large marker with standard error. The dotted lines indicate chance. **(C)** Proportion of fixation time at relevant episode locations as a function of choice (take vs. leave) and reward valence (positive vs. negative). Thin lines show individual participants; colored points show group means with standard error. **(D)** Choice prediction accuracy and coefficients from a 10-fold cross-validated logistic regression. To aggregate across trials, coefficients corresponding to each of the six encoding locations (labeled with their central degree) were used to capture how fixation time at that location interacted with the associated episode’s reward and its offer-relevance. Left: Group-level prediction accuracy with standard error across folds. Right: Regression coefficients with 95% bootstrapped confidence intervals.

We then asked whether eye gaze tracks the moment-by-moment sampling of episodic memories that support choice. If fixations reflect ongoing episodic retrieval during deliberation, then participants should direct their gaze toward the encoding locations of offer-relevant episodes. Their gaze should further be modulated by reward in a choice-dependent manner, with positively (negatively) rewarded episodes preferentially fixated before decisions to take (leave).

Consistent with these predictions, participants spent more time fixating on the encoding locations of offer-relevant episodes than those of irrelevant episodes while deliberating (*β*_0_ = 0.07, 95% *HDI* = [0.05, 0.10]; **Fig. 2B**). Further, they tended to direct their gaze toward the encoding locations of offer-relevant positively (negatively) rewarded episodes prior to taking (leaving) (*β*_decision*×*valence_ = 0.10, 95% *HDI* = [0.04, 0.16]; **Fig. 2C**; **Table S2**). No corresponding interaction was observed for offer-irrelevant episodes (*β*_decision*×*valence_ = 0.01, 95% *HDI* = [− 0.03, 0.05]), indicating that value-modulated gaze was specific to the episodes that likely contributed to each decision.

If gaze during deliberation reflects which episodes are retrieved, then these patterns should also predict participants’ choices outright. Indeed, we were able to decode people’s decisions based solely on the time they spent fixating at each encoding location (*M*_*accuracy*_ = 60.3 ± 0.9%, permutation *p* = 0.0047). Notably, only fixations to offer-relevant locations showed an effect of reward on choice (**Fig. 2D**). While this result demonstrates that fixations carry information about the upcoming decision, it does not establish that the sampled memories actively shape deliberation itself. To test this, we assumed that each fixation reflects a sample of the corresponding episode from memory. We then compared a standard drift diffusion model (DDM), in which evidence accumulates equally from all offer-relevant episodes, to an attentional DDM (aDDM) (12, 30) that weights each episode’s contribution by whether it is currently fixated (See Materials and Methods; **Fig. S2**). The aDDM provided a better account of participants’ joint choice and response time data (held out Δ*loglik*_aDDM−DDM_ = 11.3; *β*_0_ = 10.95, 95% *HDI* = [7.96, 13.91]), with unfixated locations receiving only 2.3% as much evidential weight as the currently fixated location. These findings demonstrate that eye gaze tracks episodic retrieval during choice and that samples from episodic memory, indexed by fixations, actively shape the underlying evidence accumulation process.

## Episodic sampling during deliberation is resource-rational

Having established that fixations during deliberation reflect sequential samples taken from episodic memory, we next asked *how* people sample. Specifically, because cognitive resources are limited and retrieval is costly (31–33), we examined whether deliberation is *resource-rational* with respect to the trade-off between this cost and resulting improvements to estimation accuracy. To accomplish this, we formalized deliberation as a sequential decision problem over a metalevel Markov decision process (MDP) (15, 34, 35). In our model, an agent maintains beliefs about each episode’s reward and relevance to the current offer and then uses these beliefs to estimate the offer’s value. At each time step, it either samples one of the six episodes to update its beliefs or stops deliberating and commits to a choice. Each retrieval incurs a fixed sampling cost, reflecting baseline effort, as well as a distance cost proportional to the spatial separation between successively sampled episodes, reflecting the oculomotor cost of transitioning between locations (36). An optimal agent continues sampling only as long as another retrieval is expected to refine its offer-value estimate, thereby improving its eventual choice enough to justify incurring these costs.

Identifying the optimal sampling policy is analytically intractable in the present task. We therefore trained a recurrent neural network (RNN) to approximate the optimal policy using meta-reinforcement learning, penalizing each sample with the same costs as the metalevel MDP so that it learned to balance its decision quality against retrieval effort rather than simply maximize accuracy (**Fig. 3A**) (37–40). This training yields a joint policy over which episode to sample and when to stop, producing sequences of samples (which we treat as equivalent to fixations) and choices that can be compared with human behavior. Correspondence between the network and people suggests that memory sampling is resource-rational under the same cost-benefit trade-off the network learns to solve. While the network was designed to capture retrieval during deliberation, it is not itself a model of episodic memory and therefore does not implement encoding or free recall, processes that require different architectural constraints (10, 41–44).

**Figure 3:**
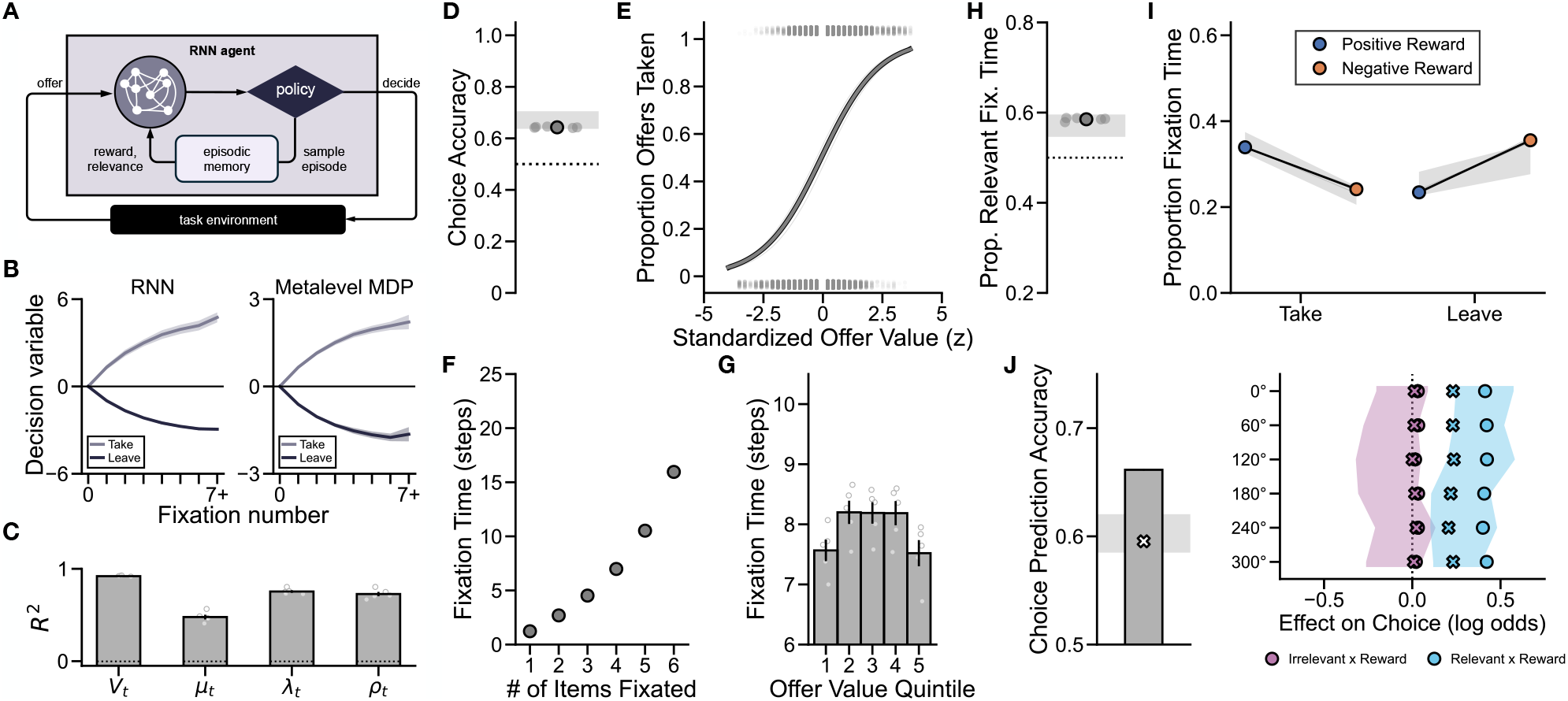
An optimal episodic sampling policy captures human behavioral and gaze patterns. A recurrent neural network (RNN) was trained to approximate a resource-rational policy. Bands show the 95% confidence interval of the human mean in panels D, H, I, and J. Across panels, individual seeds are shown as points, means across seeds as bars or larger markers, and dotted lines indicate chance. **(A)** The RNN receives an offer and selects actions at each time step. The RNN can either *deliberate* by sampling an episode from memory, which returns a noisy reward and relevance observation for the sampled episode, or *decide* to take or leave the current offer. **(B)** The RNN accumulates evidence similarly to the metalevel MDP. Left: The RNN’s decision variable (difference between the take and leave output logits) plotted as a function of fixation number and split by its eventual decision. Right: The metalevel MDP’s decision variable (*V*_*t*_ = ∑_*i*_ *µ*_*t,i*_ *ρ*_*t,i*_, where *µ*_*t,i*_ is the posterior mean reward at location *i*, and *ρ*_*t,i*_ is the posterior probability that location *i* is offer-relevant). **(C)** Cross-validated performance of a linear decoder trained to predict each component of the metalevel MDP’s belief state from the network’s hidden state at each time step. Variables are defined as in **B**, also with *λ*_*t*_, the posterior precision of the reward belief at each location. Bars show means across locations for *µ*_*t*_, *λ*_*t*_, *ρ*_*t*_; chance is estimated from permuting the outcome labels. **(D)** Choice accuracy. **(E)** Proportion of offers taken as a function of standardized offer value. **(F)** The network’s time spent deliberating (in sampling steps) as a function of the number of unique locations sampled. **(G)** Fixation time by offer value quintile, with standard error across seeds. **(H)** Proportion of fixation time directed toward offer-relevant episode locations. **(I)** Proportion of fixation time at relevant locations as a function of choice (take vs. leave) and reward valence (positive vs. negative). **(J)** Cross-validated choice prediction results. Left: prediction accuracy using all of the network’s fixations (bar) and after dropping a subset of fixations (cross). Right: Regression coefficients from the choice prediction model when all fixations were included (points) and after dropping a subset of fixations (cross). Note that panels B and C show the no prior memory network to illustrate evidence accumulation from scratch; all other panels show the prior memory network, which best matched human behavior (see below).

We first examined the trained RNN’s dynamics to verify that it learned to accumulate evidence and to represent beliefs consistent with the optimal policy in the metalevel MDP. Importantly, the metalevel MDP’s belief states are Bayesian posteriors that can be computed from the same observations received by the network, allowing us to evaluate what the network represents internally. We found that as the network sampled episodes, its decision variable progressively tracked its ultimate choice, and this closely mirrored the metalevel MDP’s estimates of offer value (**Fig. 3B**).

Each sufficient statistic of the belief state and its aggregate decision variable could also be decoded well above chance from the network’s hidden state (**Fig. 3C**; **Fig. S3**), suggesting that the network internally tracks the metalevel MDP’s beliefs and its components.

Having verified that the network approximates the optimal sampling policy, we next asked whether people deliberated in the same way by comparing the network’s behavioral and gaze signatures to those we observed in participants. Like people, the network was more likely to ‘take’ high-value offers (**Fig. 3E**) and deliberated longer both when it sampled more memories and when offers were more ambiguous (**Fig. 3F–G**). The network further captured all directly comparable behavioral and gaze effects, with no differences on any measure (**Fig. 3D,H–I**; **Table S3**) except choice prediction accuracy (**Fig. 3J**). This difference is expected, because participants’ gaze likely captures a subset of their ongoing retrieval: even during free recall, participants fixated on an episode’s encoding location only part of the time (**Fig. 2A**). To simulate this partial observability, we randomly dropped a subset of the network’s samples, retaining only the average proportion captured by participants’ gaze during free recall (42.2 ± 2.9%). This reduced the network’s choice prediction accuracy from 66.2% to 59.5%, matching people (60.3%, 95% CI = [58.5%, 62.0%]; **Fig. 3J**; **Fig. S4**).

Together, the close correspondence between human participants and an RNN that learned to make efficient choices by retrieving only when it was worth the cost of doing so suggests that human deliberation reflects similarly resource-rational sampling from episodic memory.

## Prior retrieval initializes episodic sampling

So far, our results indicate that fixations shape how evidence from episodic memory accumulates and that the way people sample this evidence resembles an optimal policy. We next sought to more fully characterize the algorithm underlying flexible decision making by first asking how this resource-rational sampler is initialized, and then what guides it from one sample to the next.

In our model, identifying which episodes are relevant to an offer can occur only by first sampling them. Initial samples are therefore unguided by memory, and preferential sampling of decision-related memories emerges only as information accrues. Yet if participants’ gaze instead reflects these preferences from the very onset of deliberation, it would suggest that retrieval precedes and guides the sampling process (e.g., via the offer cue triggering associated retrieval before deliberate sampling) rather than solely arising from it. To test this idea, we examined how these effects evolved over the course of deliberation.

We found that participants’ earliest fixations were more consistent with a sampling algorithm that was initialized by prior retrieval. We first decomposed fixations into initial visits (the first fixation to each episode’s encoding location on a given choice) and revisits (any fixation after the first). If offer-relevance is unknown at the onset of sampling, a bias toward offer-relevant locations should be absent in initial visits and emerge only with repeated sampling. Instead, these proportions were comparable (56.5 ± 1.2% vs. 56.9 ± 2.5%; **Fig. 4A**). This pattern held at nearly all ordinal fixation positions spanning the first 80% of participants’ cumulative fixation time (7.3 ± 0.9; **Fig. 4B**): participants both directed more fixations toward offer-relevant locations (*β*_0_ = 0.06, 95% HDI [0.03, 0.10]; **Fig. 4C**) and dwelled longer on them (*β*_relevance_ = 0.17, 95% HDI [0.08, 0.26]; **Fig. 4D**), consistent with active reward evaluation only for offer-relevant episodes. Reward also modulated gaze in a choice-dependent manner from the first fixation, with participants fixating more on positively (negatively) rewarded locations before deciding to take (leave) (*β*_decision*×*valence_ = 0.12, 95% HDI [0.05, 0.18]; **Fig. 4E**; **Table S4**).

**Figure 4:**
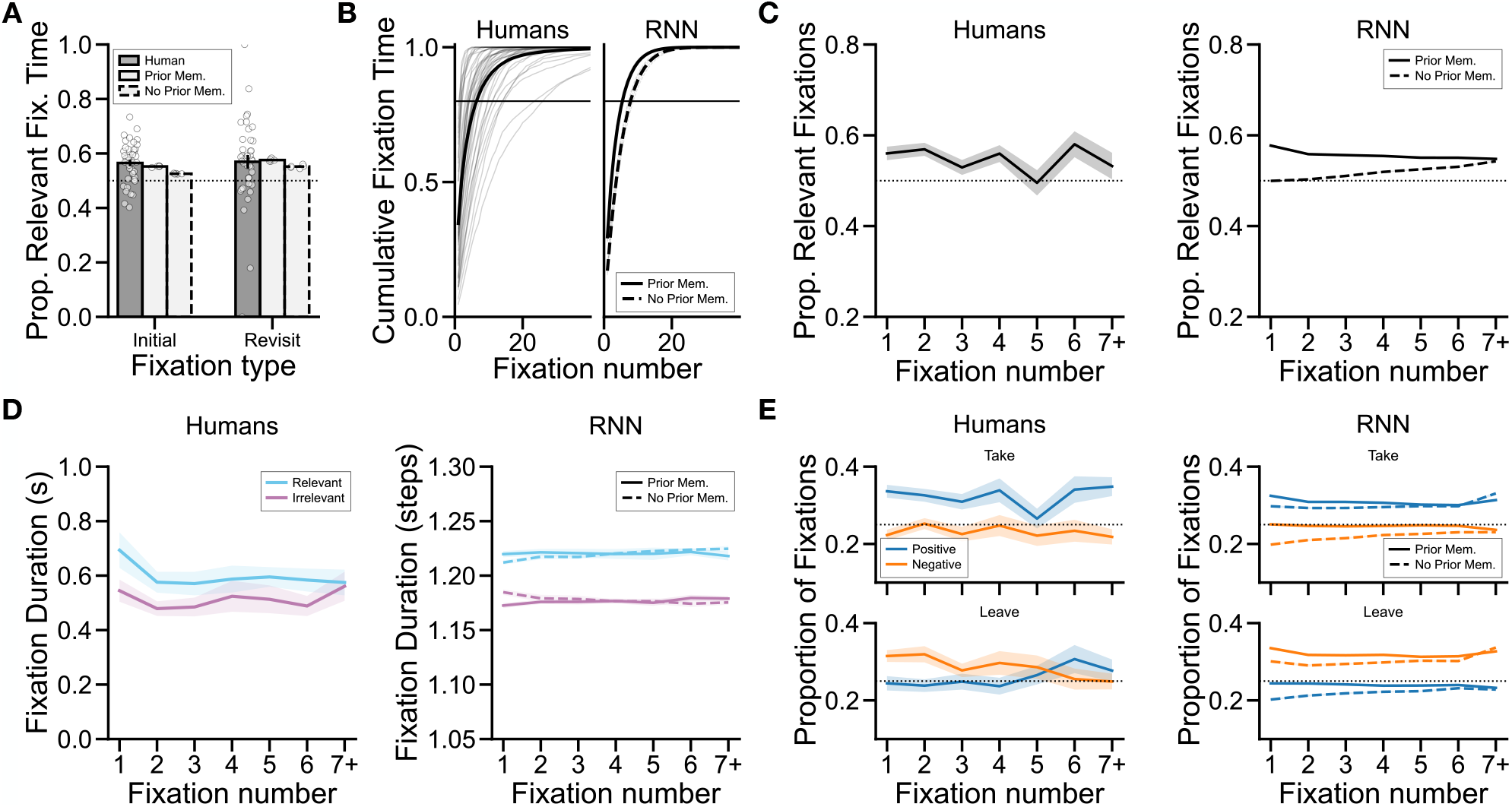
Memory guides gaze from the onset of deliberation. All panels compare human participants with the *prior memory* and *no prior memory* networks. **(A)** Proportion of fixation time on offer-relevant locations for initial visits (the first fixation to each encoding location on a given trial) versus revisits. Each point is one participant. The dotted line indicates chance. **(B)** Cumulative proportion of total fixation time as a function of ordinal fixation number. Left: Human participants, with gray lines showing individuals and the black line showing the group mean. Right: RNN variants. **(C)** Average proportion of fixations directed toward offer-relevant locations by ordinal fixation position for human participants (left) and the RNN variants (right). The shaded region shows standard error and the dotted line indicates chance. **(D)** Average fixation duration as a function of ordinal fixation position and offer relevance for human participants (left) and the RNN variants (right). Shaded regions show standard error. **(E)** Average proportion of fixations directed toward positively and negatively rewarded relevant locations, split by take (top) and leave (bottom) decisions, across ordinal fixation positions for human participants (left) and the RNN variants (right). Shaded regions show standard error and dotted lines indicate chance.

By contrast, the RNN’s early samples were unguided, with decision-related biases emerging only as it gradually accumulated information (*no prior memory* ; **Tables S3, S5**; **Fig. 4**). However, allowing it to retrieve a single noisy sample from a subset of episodes prior to deliberation eliminated every discrepancy (*prior memory* ; **Tables S3, S5–S6**; **Fig. 4**), although participants’ gaze on leave trials remained noisier at later positions (*β*_valence*×*position_ = 0.012, 95% HDI [0.004, 0.021]). With prior retrieval, the network did not have to solely discover relevance through sampling and could use this information to guide its samples from the outset. Thus, deliberation via episodic sampling is consistent with a resource-rational policy, but specifically one initialized by memories present when sampling begins.

## Rational sampling and encoding context structure deliberation

To complete our characterization of memory sampling during flexible choice, we next asked what governs the sequence of episodes people sample throughout deliberation. We considered three categories of factors: *resource-rational* effects predicted by optimal sampling, *encoding order* effects related to temporal organization in memory, and other *biases* that could plausibly influence fixations despite playing no clear role in the sampling policy. To disentangle these contributions, we modeled each fixation as arising from a choice among the possible locations using a conditional logistic regression with predictors corresponding to each factor.

Both participants and the RNN preferentially looked at locations predicted by the optimal sampling policy. Specifically, an optimal agent should stay with an offer-relevant episode only as long as each additional sample meaningfully reduces uncertainty about its reward. Once the utility of continuing to sample the current episode drops below that of sampling a less well-estimated relevant episode, the agent should switch, preferring nearby locations to minimize transition costs. Consistent with this idea, both participants and the network were more likely to fixate on the locations of offer-relevant episodes that had received less cumulative fixation time (inverse relevant fixation time: *β*_human_ = 0.061, 95% HDI = [0.033, 0.089]; *β*_network_ = 0.115, 95% HDI = [0.112, 0.118]; **Fig. 5A**; null comparison in **Fig. S5**). Likewise, each new fixation was also more likely to land on locations that were physically closer to the previously fixated location (humans: *β*_cw_ = −0.069, 95% HDI = [−0.101, −0.039]; *β*_ccw_ = −0.108, 95% HDI = [−0.151, −0.066]; network: *β*_cw_ = −0.045, 95% HDI = [−0.048, −0.042]; *β*_ccw_ = −0.040, 95% HDI = [−0.043, −0.037]). Notably, participants were more sensitive overall to spatial proximity than the network (*β*_human_ = 0.28, 95% HDI = [0.24, 0.33] vs. *β*_network_ = 0.12, 95% = HDI [0.10, 0.15]; **Fig. 5B**; **Fig. S6**; **Table S8**), possibly reflecting additional effects of spatial encoding context on memory retrieval (45–47) beyond the oculomotor cost captured by our model.

**Figure 5:**
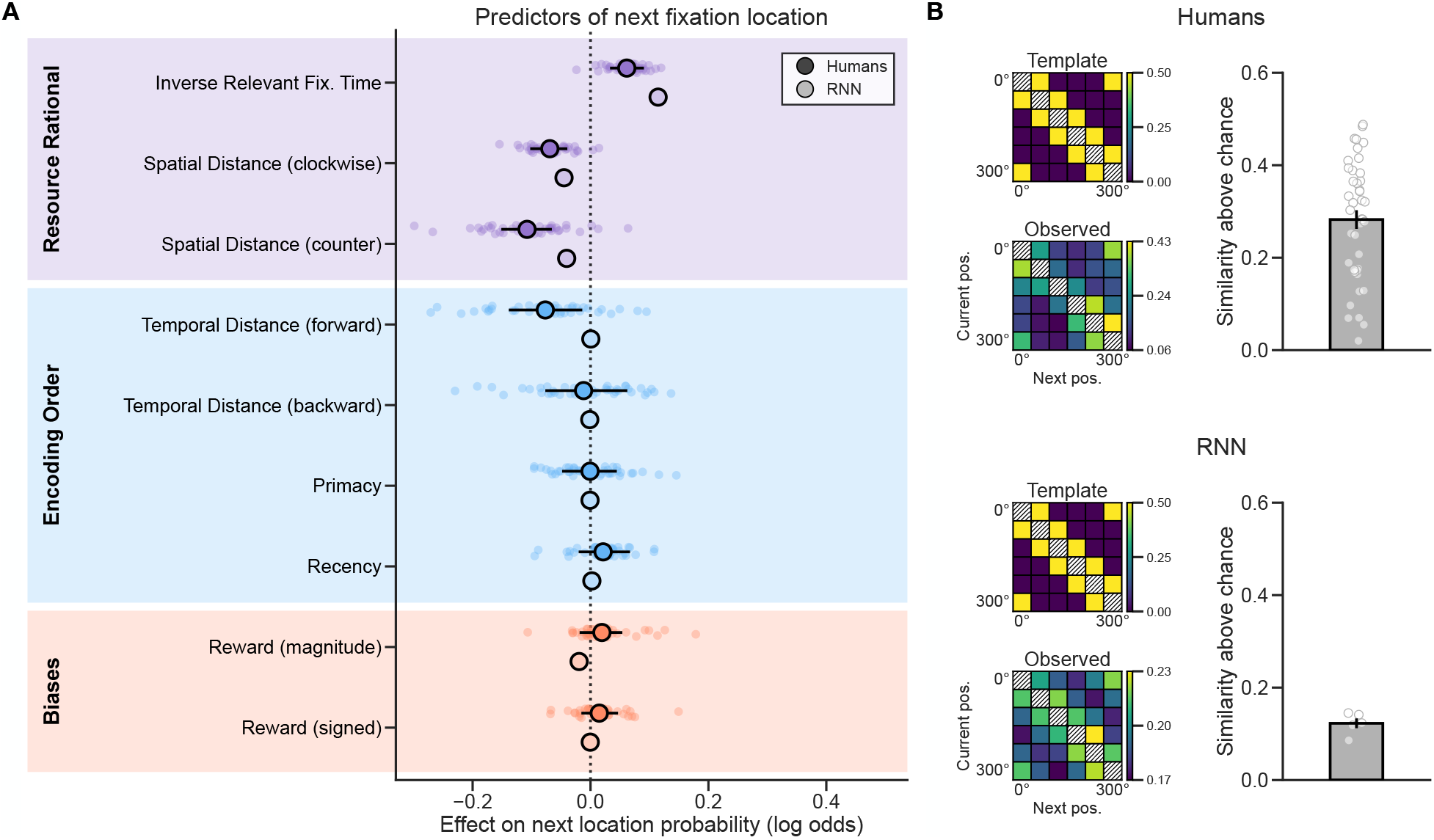
Determinants of memory sampling throughout deliberation. **(A)** Regressors (z-scored) organized into three categories were used to predict the location of each new fixation during deliberation. Points and horizontal lines indicate the group-level posterior mean and 95% HDI for humans (dark) with small points showing per-subject random effects and for the prior memory network (light). Predictor categories are denoted by background color. The dotted line at zero indicates no effect. See Table S7 for all group-level estimates. **(B)** Spatial fixation effects. Left: Fixation transition structure. A template transition matrix showing the expected pattern when fixations move between spatially adjacent encoding locations, alongside the observed transition matrix computed from actual fixations. Right: Pearson correlation between the template and observed matrices relative to permuted chance. Error bars represent standard error and points are one participant or network seed.

How information is temporally organized at encoding is also known to shape retrieval: items at the start and end of study are recalled more readily (primacy and recency), and items studied close in time tend to be recalled together (41, 42). We thus asked whether organizational effects of this kind surface during deliberation, even though the optimal policy is indifferent to them. Indeed, participants were more likely to fixate on the locations of episodes encoded close in time to the episode at the location that was just fixated, with this effect concentrated in the forward direction (forward temporal distance: *β*_human_ = − 0.077, 95% HDI = [− 0.140, −0.015]; **Fig. 5A**). Primacy and recency had no detectable effect. Unsurprisingly, the network showed no influence of encoding order on any measure. This contrast demonstrates that human memory sampling during deliberation reflects not only the optimal policy, but also the temporal structure of memory.

Taken together, these findings indicate that retrieval during deliberation is jointly shaped by optimal sampling and the organization of memory itself, with no substantive evidence for biases not predicted by either of these factors (see supplementary text).

## Discussion

We found that as people deliberated, they directed their gaze toward the encoding locations of decision-relevant episodes. Their fixations tracked both the content and dynamics of retrieval, and an RNN trained to approximate an optimal sampling policy reproduced these patterns. Our results provide direct evidence that humans take samples from episodic memory to construct ad hoc decision variables and identify a resource-rational algorithm that governs deliberation: at each moment, sampling targets memories that meaningfully reduce decision uncertainty and halts when expected gains in decision quality no longer outweigh retrieval costs. This sampler is initialized by prior retrieval and biased by how memory is organized at encoding.

Prior work on value-based choice has shown that gaze modulates evidence accumulation (12, 30), even when options are not visible (48, 49). Yet decisions in these tasks are typically between prescribed items with known values, and fixations index which option is currently attended. Comparatively little is known about how people deliberate when value must instead be built from memory. Here we show that when value is assembled over multiple episodes, gaze to each encoding location reveals both which memories are sampled and how they accumulate toward a decision.

A remaining question concerns the role of gaze itself in deliberation. Whether fixations actively facilitate retrieval by reinstating the spatial encoding context (28, 50) or whether they instead capture “leakage” from a retrieval process already underway (51) is unclear. Indeed, spontaneous motor behavior may reflect latent cognitive states (52), and fixations during deliberation could similarly signal retrieval. Our findings suggest that retrieval at least partly directs gaze: fixations were biased toward offer-relevant episodes before any overt sampling had occurred, and the network reproduced this pattern only when initialized with prior samples. What determines which episodes are available before deliberation is an open question. Whether gaze also feeds back to guide retrieval can be tested in future work by, for instance, restricting gaze during deliberation, as has been done in explicit recall tasks (50, 53).

Our model treats episodic memory as a fixed store from which noisy samples are drawn but does not account for how episodes are initially encoded. Yet how information is organized at encoding shapes how it is later retrieved, and during deliberation participants preferentially fixated the encoding locations of episodes studied just after the episode whose location was currently fixated. This forward bias is a hallmark of temporal context theory (41, 42). A similar pattern emerged spatially: although both people and the network tended to fixate on locations that were physically close to one another, participants weighed spatial proximity more strongly. Given the well-established role of spatial context in organizing episodic memory (45–47, 54), this finding suggests that the spatial layout at encoding likely shaped retrieval in ways not fully captured by the network. Integrating our model of deliberation with RNN architectures that explicitly model encoding and retrieval mechanisms (41–44) thus remains an important research direction to account for these effects.

Relatedly, a full neural account of memory-guided choice must capture both how the brain stores memories as well as how they are sampled and combined into a decision. Encoding and retrieval mechanisms are likely supported by the hippocampus (41, 44), whereas the deliberative control that selects which memories to sample may depend on the prefrontal cortex (55). Our approach speaks to this prefrontal contribution. Specifically, our RNN was trained through meta-reinforcement learning, which has been proposed as an explanation for how the prefrontal cortex learns to efficiently deploy flexible reward-guided behaviors (38). The recurrent dynamics it acquires thus offer a candidate account of how prefrontal circuits might carry out resource-rational sampling during deliberation. This possibility is bolstered by recent work showing that networks trained to select mental computations in this way capture features of prefrontal neural activity (40).

Finally, the use of gaze as a readout of memory-guided deliberation opens a new methodological path for studying retrieval dynamics during choice. This has remained largely out of reach in humans. Although neuroimaging has traditionally been used to measure memory reactivation during choice (17, 56), deliberation unfolds over extended and variable timescales, making it difficult to determine when retrieval actually occurs. Using fixations to capture retrieval as it unfolds solves this problem, and recent work has shown that gaze corresponds to memory reactivation at the neural level (57, 58). More broadly, because this approach requires only that memories be encoded in distinct spatial locations, it should be applicable to any task in which adaptive behavior relies on associative memory.

Together, these results demonstrate that flexible decisions are constructed through sequential sampling from episodic memory, and that this process is governed by the same resource-rational principles that structure simpler forms of choice and, arguably, cognition in general.

## Materials and Methods

### Participants

We recruited students and New York University affiliated community members from the NYU subject pool. Participants were paid a base rate of $15/hour with the opportunity to earn up to $15 in bonus money. We recruited a total of 50 participants (30 female, 15 male, 5 other; *M*_*age*_ = 22.07 ± 2.78 years, range = 18.2 − 31.2) with normal or corrected-to-normal (via contacts) vision. Based on pre-registered criteria (59), participants were excluded if an adequate 13-point eye-tracking calibration was not achieved at the beginning of the study (3) or if they achieved less than 70% accuracy on a pre-task comprehension quiz (4), leading to a final sample of 43.

### Experimental Procedure

Participants completed a four-part task based on a recent behavioral study (1). Completing all four parts (a “round”; note that we found no effects of round number of any behavioral or eyetracking result; **Table S9**) took approximately five minutes, and participants completed seven rounds. Participants’ eye gaze was recorded monocularly at 1000 Hz throughout the entirety of each round using an EyeLink 1000 Plus with a chin rest (SR Research Ltd., 2013). Each round began with a drift check, and we recalibrated the eye tracker if the drift check failed.

### Encoding Phase

In the first part of a round (**Fig. 1A**), participants completed a task designed to allow them to encode 6 individual items and an associated reward (an “episode”). Each item was presented on the screen for one second, after which its reward appeared alongside it for another six seconds. An item’s associated reward consisted of a pseudo-randomly sampled integer (excluding 0) between either -9 and 9. Each item had four different binary features: texture (a solid color or with a pattern), environment (land or sea), animacy (animal or object), and size (larger or smaller than a microwave). Immediately after viewing each item and its associated reward, participants completed an attention check consisting of the item alongside two options, either the associated reward that was just shown or another randomly selected reward. They had three seconds to respond by pressing either the F key if the correct reward appeared on the left, or the J key if it appeared on the right. Each item and reward was viewed only once per round, for a total of six encoding trials per round. Importantly, each item and its reward was presented in one of six possible locations evenly distributed around a circle. Items could repeat across rounds, and the number of repetitions per item was pseudo-randomly balanced within each session. Each time an item appeared, it was associated with a different reward, but it always appeared in the same location.

### Distractor Phase

In the second part of a round, participants completed a 90 second distractor task to prevent active rehearsal of the episodes. This distractor consisted of a 2-back working memory task in which participants were shown one of several letters in sequence. Participants were asked to identify whether the current letter matched the one presented two steps earlier by pressing the F key.

### Decision Phase

In the third part of a round, participants made eight decisions based on each of the features that were seen in a round (animal/object, large/small, land/sea, solid/pattern). Each decision consisted of an offer in which a single feature (e.g. animal) was said out loud, and participants were then asked to either take or leave this offer. Participants were informed that the value of each offer consisted of the sum of each episode that was described by the offer (e.g. the value of the animal offer would be the sum of the rewards associated with all animals seen during encoding), and that they should take positive and leave negative offers. Participants had an unlimited amount of time to make each decision and used the F key to take the offer and the J key to leave the offer. Importantly, during decision making the screen was entirely blank except for the circle around which each item and its associated reward appeared during encoding. Note that we intentionally selected features that differed in their overall saliency (i.e. animacy versus texture) to reproduce the variety of features that appear in real-life decision making, and we found that these differences do not systematically impact any results (**Fig. S7**).

### Memory Phase

In the fourth part of a round, participants completed a memory phase in which there were three memory tests: a free recall test, a reward recall test, and a location recall test. First, following a brief auditory stimulus signaling the start of the trial, participants were asked to freely recall out loud the names of each of the items they could remember seeing during the current round. During the reward recall test, participants then heard the name of each item that was shown in a round, and they were asked to recall its associated reward out loud before pressing a key to move on to the next item. Finally, during the location recall test, participants again heard the name of each item out loud, but were asked to select which of the six possible positions it appeared in.

### Round Generation Procedure

The items and associated rewards of each round were generated according to the following pseudo-random procedure. First, combining across the four binary feature dimensions yielded 16 unique items. In each round, six of these 16 items were selected such that exactly three items were of each of the two possible types for every feature dimension (e.g., three animals and three objects), ensuring that every decision involved exactly three offer-relevant and three offer-irrelevant items. Item reward values were constrained so that the offer-relevant items for each decision summed to a non-zero value and contained a mixture of positive and negative rewards. The full configuration of items and rewards across all seven rounds for each participant was then selected from 1000 randomly generated candidate configurations via a scoring procedure that optimized for balanced item and position usage across rounds, minimized correlation between summed relevant and irrelevant reward values, and allowed for an equal number of take and leave decisions, as well as an equal number of items with positive and negative associated rewards across all decisions.

### Pre-task Training

Prior to completing the primary task, participants completed a 10 minute training designed to teach them the locations that each image appeared in. Image-location mappings were pseudo-randomly assigned across participants. In this training, participants completed six blocks in which they were first shown the locations of six out of sixteen total images. Each image’s label was then read out loud three times per block, and participants were required to respond with the location of that image. Following these blocks, in a final session participants were then read each of the sixteen images labels out loud and were required to respond with the image’s location. Each image was repeated twice, and participants were required to respond accurately both times. If they missed one of the responses, this trial was added to the end of the procedure.

### Behavioral Analysis

Unless otherwise specified, all data were analyzed with regression models estimated using hierarchical Bayesian inference such that group-level priors were used to regularize participant-level estimates. All predictors were specified as fixed effects alongside random slopes and intercepts that were allowed to vary across subjects. The joint posterior was approximated using No-U-Turn Sampling as implemented in stan (60). Four chains with 2000 samples (1000 discarded as burn-in) were run for a total of 4000 posterior samples per model. Chain convergence was determined by ensuring that the Gelman-Rubin statistic R was close to 1. Default weakly-informative priors implemented in the brms package were used for each regression model (61). For all models, fixed effects are reported in the text as the mean of each parameter’s marginal posterior distribution alongside 95% highest density intervals (HDIs), which indicate where that percentage of the posterior density falls. Parameter values outside of this range are unlikely given the model, data, and priors. Thus, if the range of likely values does not include zero, we conclude that a meaningful effect was observed.

### Decision Analysis

We first assessed overall performance in the decision phase using a mixed-effects logistic regression model. The model included fixed (*β*) and random (*u*) intercepts to determine whether accuracy (correct = 1, incorrect = 0) was different from chance-level performance:

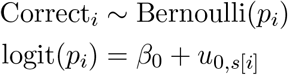

where *s*[*i*] indexes the participant for observation *i*.

We next asked whether choices were better predicted by an offer’s true value or by participants’ recalled offer value. The true offer value for a given trial was defined as the sum of rewards associated with all items matching the offered feature, whereas the recalled offer value was defined as the sum of recalled rewards for the subset of offer-relevant items that participants successfully remembered during the free recall task. To facilitate comparison, both predictors were *z*-scored prior to model fitting. We fit two logistic mixed-effects models predicting choice (take = 1, leave = 0) with random intercepts and random slopes per participant. The true offer value model was specified as:

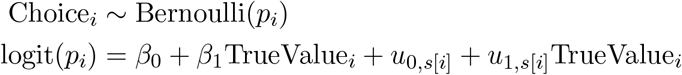

and the recalled offer value model was:

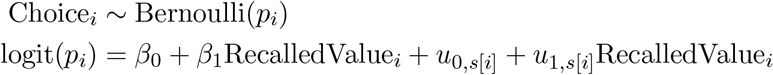

where TrueValue and RecalledValue denote *z*-scored true and recalled offer values, respectively.

#### Model comparison

To determine the extent to which either recalled offer value or true offer value best predicted choices, we compared these models using 10-fold cross validation. The expected log pointwise predictive density (ELPD) was then computed by summing the log likelihood for each held out datapoint and used as a measure of out-of-sample predictive fit for each model. Higher ELPD values suggest better model fit, as they indicate a higher likelihood of accurately predicting new data. To compare models, we then subtracted the pointwise ELPD estimates and calculated the standard error of this difference to quantify uncertainty of the comparison(62, 63).

### Response Time Analysis

To determine whether decision response time (*RT*) varied as a function of offer value, we fit two mixed-effects models predicting the log-transformed response time of each decision using linear and quadratic terms with both random intercepts and random slopes per participant:

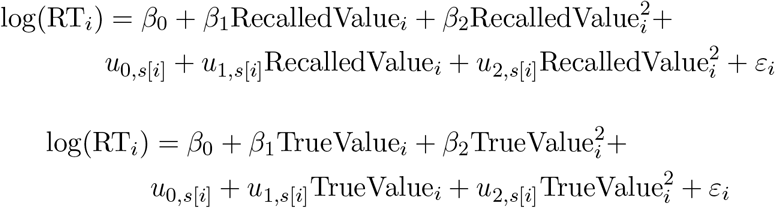

where *ε*_*i*_ ∼ *N* (0, *σ*^2^).

To next examine how the number of recalled memories impacted the amount of time it took participants to make their choices, we used a linear mixed-effects model to predict trial-wise decision response times from the total number of memories participants recalled in each round (nMemories):

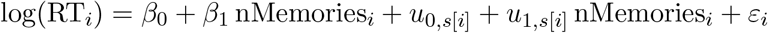

### Memory Analysis

First, to assess overall recall performance, we computed the proportion of the six encoded items that each participant freely recalled in each round and compared these recall rates with chance performance, which we defined as the probability of correctly guessing items when randomly selecting 6 items from the pool of 16 possible items that could be shown on a given round, which was 37.5% (or 6/16). For each round, we then computed the difference between the mean recall rate and chance (*δ*_recall,*i*_) and then fit a mixed effects linear regression model to these difference scores:

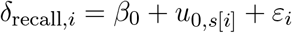

Next, to determine the accuracy of reward memory, we regressed the true reward of each item on the reward recalled by each participant:

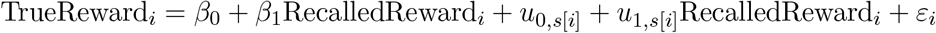

with random intercepts and random slopes per person.

Lastly, we tested whether performance on the spatial memory task exceeded the chance rate of 1*/*6 (one of six possible locations). We first computed the proportion of the six item locations that each participant recalled correctly in each round, subtracted chance (*δ*_spatial,*i*_), and then fit a mixed effects linear regression model:

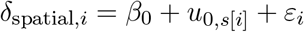

### Eye-Tracking Analysis

As outlined in our preregistered criteria (59), individual trials were excluded from eyetracking analyses if greater than 50% of the eyetracking data was lost (e.g. due to blinks or loss of eye position). Two individual participants were excluded from eyetracking analyses entirely because greater than 50% of their trials were rendered invalid by the previous criterion, yielding a final sample of *n* = 41 for all eyetracking results.

All eyetracking analyses focused on fixations (defined using the EyeLink online parser) during the decision phase and free recall task. Areas of interest (AOI) for each item were defined as the segment of the circle shown on the screen that it appeared within during the encoding phase. Each AOI consisted of 530 pixels plus a 50 pixel buffer to account for measurement noise. A fixation was assigned to an AOI if the gaze position fell within these pixels.

### Free Recall

Fixations during free recall were analyzed by examining the proportion of fixation duration allocated to the encoded location of each recalled item. We identified the timepoint at which verbal recall of an individual item was initiated, extracted a window (from -3000 to +750 ms) around this timepoint, and calculated the duration spent fixating at the AOI of the recalled item divided by the total fixation duration across all image AOIs within 100 ms segments of the overall timeseries. These fixation proportions were computed separately for each recall event and then averaged within participants.

We then assessed statistical significance using a cluster-based permutation test. At each time bin, we computed each participant’s deviation from the chance level of 1*/*6 (i.e., equal looking across six locations). We then performed 1000 sign-flip permutations of these subject-level deviations. Adjacent time bins exceeding a cluster-forming threshold of *p <* 0.01 were grouped into clusters, and the summed test statistic within each cluster was compared to the null distribution of maximum cluster statistics. Clusters were considered significant at *α* = 0.05, with a minimum cluster size of 8 time bins.

### Decision Phase

Fixations during the decision phase were analyzed by first classifying item locations as *offer-relevant* (associated item matched the offered feature; which was always 3 of 6 items) or *offer-irrelevant* (associated item did not match the offered feature; which was the remaining 3 items). Item valence (positive or negative) was determined using participants’ recalled reward values when available, with a fallback to the true encoded values in the rare occasion that the participant did not give a response for an item’s reward due to accidental button presses. Consecutive fixations that were directed to the same AOI were merged prior to analysis. Unless otherwise specified, all analyses consisted of Bayesian mixed-effects regression models, using the same estimation settings as the behavioral models (4 chains, 2000 iterations with 1000 burn-in per chain).

#### Fixation time by offer relevance

We first asked whether fixation time to offer-relevant locations exceeded chance (50%), which we captured as the trial-level deviation of the proportion of fixation time to relevant locations from chance:

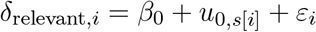

where *δ*_relevant,*i*_ is the proportion of fixation time directed to offer-relevant locations minus chance on trial *i*.

#### Fixation time by valence and decision

Next, we examined the proportion of fixation duration allocated to positively-valenced and negatively-valanced offer-relevant item locations during each choice. We computed the proportion of time spent fixating on the AOIs corresponding to each item’s encoding location by summing fixation durations to each AOI and dividing by the total across all six item AOIs in the trial. We then modeled deviation from chance as a function of the decision (take vs. leave) and valence label (positive vs. negative). Chance for each trial was computed as the proportion of positive/negative items present (between 1 and 3) divided by the total number of items (6). We then fit the following mixed effects model:

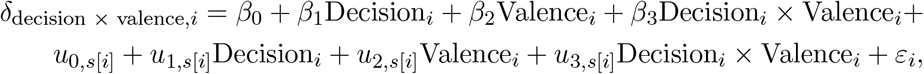

where *δ*_decision *×* valence,*i*_ is the proportion of fixation time directed to offer-relevant positive or negative locations minus chance on trial *i*, and Decision_*i*_ and Valence_*i*_ are indicators for decision type and valence, respectively. An analogous model was fit using instead the raw fixation proportion as the dependent variable, and the same set of models was fit to offer-irrelevant item locations to assess whether the decision- and valence-dependent gaze patterns observed for offer-relevant items extended to irrelevant items.

### Predicting choice from fixation patterns

We also assessed whether fixations during decision making were explicitly predictive of participants’ choices. Doing so requires representing fixations across multiple trials using a set of shared coefficients. To accomplish this, we took advantage of the incidental encoding locations of each episode. Specifically, at the group-level we fit an *L*_2_-regularized logistic regression model predicting binary choices (take = 1, leave = 0) from features encoding the interaction between gaze allocation and an item’s relevance and reward at each encoding location around the circle. For each trial, we defined 18 features drawn from the six encoding locations (*i* = 1, …, 6). The feature vector for each trial consisted of:

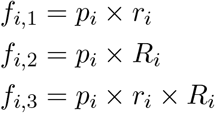

where *p*_*i*_ denotes the proportion of total fixation time directed to location *i, r*_*i*_ denotes the recalled reward value of the item at location *i*, and *R*_*i*_ ∈ {0, 1} denotes whether the item at location *i* was offer-relevant. Because *R*_*i*_ was coded as 0 and 1, regression coefficients fit to the first set capture gaze weighted by the interaction between irrelevant items and reward, those fit to the second set capture gaze as it is impacted by relevance, and those fit to the third capture gaze weighted by the interaction between relevant items and reward. Model performance was evaluated via 10-fold cross-validation. Specifically, within each fold, features were *z*-scored using the mean and standard deviation of the training set, and classification accuracy was computed as the proportion of correctly predicted choices across held-out folds. To estimate uncertainty around model coefficients, we performed 1000 bootstrap resamples and report 95% confidence intervals derived from the bootstrap distribution of each coefficient.

To ensure that gaze added useful predictive signal beyond what could be obtained from chance, we constructed a permutation null distribution. For each of 10000 permutations, we independently shuffled the assignment of items to encoding locations within every trial. For each permutation we recomputed the 18 features and reran the same 10-fold cross-validation procedure. We report the one-tailed permutation *p*-value, defined as the proportion of permuted accuracies that met or exceeded the observed accuracy across all 10000 permutations.

#### Evaluating how fixations evolve over the course of deliberation

We further conducted several analyses to examine how fixations evolved over the course of deliberation. First, we asked how duration evolved over time by fitting a linear mixed-effects model predicting the log-transformed duration of each fixation from a binary indicator of whether the fixated location was offer-relevant (*R*_*i*_), the ordinal fixation position within a trial (Pos_*i*_; positions 1–6 individually, with 7+ collapsed in a single bin to account for data sparsity beyond this position), and their interaction, with random intercepts and slopes for all predictors per participant:

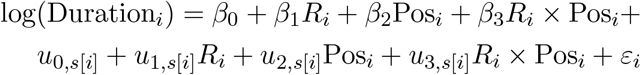

Next, to assess whether the tendency to fixate on offer-relevant locations was present throughout deliberation, we aggregated fixations within each participant and position and computed the proportion of fixations directed to offer-relevant locations minus chance (0.5). We then fit a linear mixed-effects model with position as a fixed effect and random intercepts and slopes per participant:

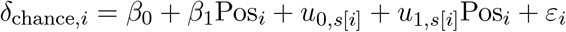

A positive intercept indicates that fixations were biased toward offer-relevant locations above chance, and the slope captures whether this bias changed across fixation positions.

We also asked whether the valence of recalled rewards modulated gaze allocation differently depending on the participant’s decision at each fixation position. For each trial and ordinal fixation position, we computed the proportion of fixations directed to offer-relevant items with positive and negative recalled rewards and subtracted the corresponding chance proportion (number of items in that category divided by six). We then fit a three-way linear mixed-effects model with decision type (Dec; take = +0.5, leave = − 0.5), valence (Val; positive = +0.5, negative = − 0.5), and position as fixed effects, with random intercepts and slopes for decision, valence, and position per participant:

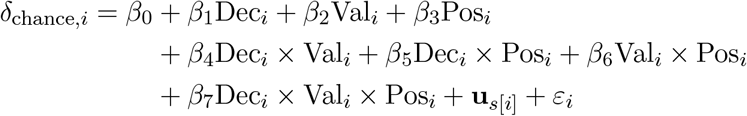

where **u**_*s*[*i*]_ includes random intercepts and random slopes for decision, valence, and position per participant.

## Attentional Drift Diffusion Model

We further modeled participants’ take/leave decisions using the attentional drift diffusion model (aDDM) (12, 30), which we altered for the present task. The model extends the standard drift diffusion model (DDM) by modulating evidence accumulation according to a participant’s current fixation location, allowing us to test whether gaze-contingent weighting of item-associated reward values improved the joint prediction of choice and response times.

### Model Specification

On each trial, the six item locations are indexed *i* = 1, …, 6, each with an associated item. Each item has a reward *r*_*i*_ and a binary relevance indicator *R*_*i*_ ∈ {0, 1}, where *R*_*i*_ = 1 for the three offer-relevant items and *R*_*i*_ = 0 for the three offer-irrelevant items. The model tracks a relative decision variable *V*_*t*_, initialized at *V*_0_ = 0, that accumulates evidence over time in favor of taking (*V*_*t*_ *>* 0) or leaving (*V*_*t*_ *<* 0) the offer. The model has three free parameters: a drift scaling factor *d >* 0, an attentional discount *θ* ∈ [0, 1] applied to unfixated offer-relevant item locations, and a diffusion noise standard deviation *σ >* 0.

At each time step *t*, the fixated item location *j* determines the attentional weights assigned to each item. When the fixated location is associated with an offer-relevant item (*R*_*j*_ = 1), the weights are

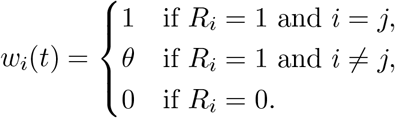

Note that *θ* has a direct interpretation as the fraction of evidential weight contributed by each unfixated relevant item relative to the fixated one. The attended offer value at time *t* is:

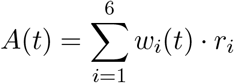

and evidence accumulates in discrete time steps of 1 ms:

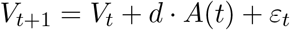

where *ε*_*t*_ ∼ *N* (0, *σ*^2^) is Gaussian noise sampled independently. Note that when the fixated item location is offer-irrelevant (*R*_*j*_ = 0), this leads evidence accumulation to pause because the drift term is set to zero while noise continues to perturb the relative decision variable. The process terminates when *V*_*t*_ reaches one of two fixed absorbing boundaries: *V* = +1 (take) or *V* = −1 (leave). When *θ* = 1, all relevant items receive equal weight regardless of which item is currently fixated, and the model reduces to a standard DDM with drift rate proportional to the sum of offer-relevant rewards. We used participants’ recalled reward values for *r*_*i*_. As in prior analyses, when a recalled value was unavailable for a given item, its true reward was used as a fallback.

### Gaze Generation Procedure

As in prior work (12), we fit the model to group-level data using simulated trials, which requires defining a generative model of gaze. Following (12), we estimated gaze statistics directly from the empirical fixation data, conditioned on recalled offer value *V*_offer_, binned into seven quantile bins. Each simulated fixation was generated IID, conditional on the *V*_offer_ bin, using the following procedure:

1. Sample whether the fixation targets a relevant or irrelevant item location with probability *P* (relevant | *V*_offer_), estimated from the empirical data.
2. Sample the fixation duration from the relevance-specific empirical duration distribution for that *V*_offer_.
3. Sample an item location uniformly from among the three relevant or irrelevant items present on a trial, with the constraint that the same item location could not be sampled twice in a row.

As in (12), we also sampled the total transition time (i.e. the sum of inter-fixation gaps within a trial) once per simulated trial from the empirical distribution of transition times and added this to the simulated decision time in order to produce a predicted response time.

### Simulation-Based Likelihood

We estimated the likelihood of the observed data under candidate parameters using a simulation-based approach similar to (12). Within each combination of *V*_offer_ bin and decision type (take/leave), observed response times were binned into 15 quantiles, with a minimum of 25 trials per bin.

For a candidate parameter set (*d, θ, σ*), we simulated 1000 trials per *V*_offer_ bin using the generative gaze procedure. Each simulated trial produced a choice and a response time which we assigned to the appropriate *V*_offer_, choice, and RT bin. To avoid log(0) from sparse bins, we computed predicted bin probabilities using Dirichlet smoothing:

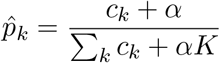

where *c*_*k*_ is the number of simulated trials falling in bin *k, K* is the number of bins, and *α* = 1.0. The log-likelihood was then computed as

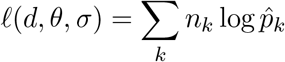

where *n*_*k*_ is the observed count in bin *k*.

### Modeling Fitting and Comparison

As in (12), during fitting *σ* was reparameterized as *µ* = *σ/d*, such that *σ* = *d*· *µ*. This reparameterization reduces the correlation between *d* and *σ* in the likelihood surface, improving optimizer performance. Parameters were estimated by maximizing the simulation-based log-likelihood using pyBADS (Bayesian Adaptive Direct Search), with parameter bounds defined as *d* ∈ [10^−5^, 10^−3^], *θ* ∈ [0.01, 0.99], and *µ* ∈ [1, 100]. We then assessed each model using 10-fold cross-validation. To mitigate sensitivity to optimizer initialization, the fitting procedure was repeated with five random seeds, and the seed yielding the highest mean held-out log-likelihood across folds was selected. Parameters are reported as the mean parameter estimate across all folds.

To examine the extent to which gaze impacted participants’ choices and response times, we compared the aDDM with an identical model but with *θ* fixed at 1. This is equivalent to the standard DDM because all associated item rewards contibute equally. Both models were fit with identical fold assignments ensuring that the same held-out trials were used to evaluate each model. For each fold *n*, we computed the held-out log-likelihood 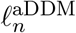 and 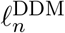, and then the fold-level difference 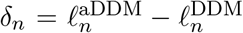 . To assess whether *δ*_*n*_ was reliably different from zero, we fit an intercept-only Bayesian linear regression model using the same estimation settings as all other models (4 chains, 2000 iterations with 1000 burn-in per chain):

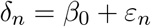

### Parameter Recovery

Lastly, to validate the identifiability of model parameters, we performed a parameter recovery analysis. We sampled 1000 parameter combinations of the three free parameters. For each combination, we generated a synthetic dataset by simulating one trial for every actual trial completed by participants using the generative gaze procedure and candidate parameters. Each synthetic dataset was then refit using the procedure described above. We then compared the correlation between true and recovered parameter values to assess recovery quality.

### Episodic memory sampling as a metalevel Markov decision process

In addition to the above analyses, we aimed to determine the extent to which people sampled optimally from episodic memory during deliberation. To this end, we formalized optimal memory sampling in the present task as a metalevel Markov decision process (MDP) (15, 34, 35). As in a standard MDP, a metalevel MDP is defined by a set of states (a set of beliefs), a set of actions (mental computations), a transition function, and a reward function. Our model is based on a metalevel MDP originally developed for simple choice tasks (15) and adapted to the present task.

### Model Specification

We first assume that each fixation to an episode’s encoding location is a sample from episodic memory of that episode’s relevance to the current offer and its associated reward. Belief states *b* ∈ ℬ in our task thus correspond to posterior distributions over the six rewards associated with each episode in a trial as well as posterior beliefs about which episodes are offer-relevant. We assume that associated reward posteriors are Gaussian, allowing the reward belief to be represented by two vectors, ***µ*** and ***λ***, specifying the mean and precision of each episode’s reward distribution. The belief additionally contain a vector ***ρ*** where *ρ*^(*i*)^ ∈ [0, 1] denotes the current posterior probability that episode *i* matches the current offer *f* . Beliefs also encode the episode sampled last to model distance-dependent costs, as well as the current offer *f* :

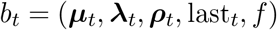

where last_0_ = ⊘ at trial onset.

A computation *c* ∈ *C* corresponds to sampling one episode (*c* ∈ {1, …, 6}) and updating beliefs, or terminating sampling and deciding, ⊥. Given belief state *b*_*t*_ and cognitive operation *c* ∈ {1, …, 6}, the next belief state *b*_*t*+1_ is generated by drawing a noisy memory sample

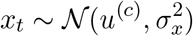

and updating the posterior reward for episode *c* according to the Bayes rule. Upon sampling episode *c* the agent also receives a feature observation with probability *k*:

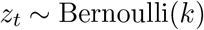

If *z*_*t*_ = 1, the agent observes the true indicator *s*^(*c*)^ ∈ {0, 1} of whether episode *c* is offer-relevant and updates *ρ*^(*c*)^. If *z*_*t*_ = 0, no feature information is obtained and *ρ*^(*c*)^ is left unchanged. All remaining items’ beliefs are unchanged and last_*t*+1_ = *c*. Note that for actual implementation (see below), we assume that *k* = 1, meaning that each fixation deterministically reveals the sampled episode’s relevance. Two empirical observations support this assumption. First, participants’ choices were well predicted by offer-relevant items but showed no influence of offer-irrelevant items (**Fig. 2D**), suggesting that participants reliably distinguished relevant from irrelevant episodes. Second, the proportion of fixation time directed toward offer-relevant locations was stable from the first fixation onward (**Fig. 4**), indicating that relevance was not learned over the course of deliberation. These findings suggest that participants had little uncertainty about episode relevance.

As in (15), we define a metalevel reward function that incorporates the cost of computation and the utility of an action. We define the arc distance between two episodes on the circle of *N* = 6 episodes as the shortest length:

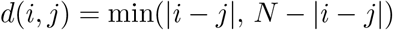

The metalevel reward for sampling from memory is:

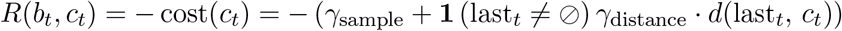

which states that there is a fixed cost for retrieving a sample from memory, *γ*_sample_, as well as a distance cost proportional to *d*(last_*t*_, *c*_*t*_), the arc distance between the previously sampled episode and the currently sampled episode, scaled by *γ*_distance_. Note that when *c*_*t*_ = last_*t*_, the distance is zero and no additional cost is incurred beyond the base sample cost.

Finally, the metalevel reward for terminating sampling is the expected utility of the optimal take/leave decision given the current belief state:

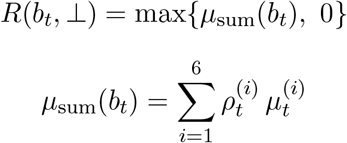

Thus, at each time step the agent chooses which episode to sample information from in its episodic memory or whether to terminate sampling and decide, trading off the expected improvement in utility against the cost of additional recall.

### Optimal metalevel policy

We assume that decisions about where to fixate (i.e. which memory to sample) and when to stop sampling are made optimally, subject to the costs specified above. We define this as the optimal policy over computations.

Let *T* denote the termination time (the first time step such that *c*_*T*_ =⊥). Terminating sampling at time *T* induces an external decision

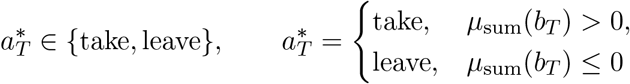

The total payoff for a single decision is the termination reward minus the cumulative cognitive costs of sampling:

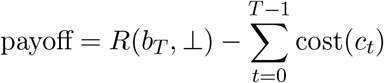

An optimal metalevel policy *π*^*^ is a Markov policy that selects computations *c*_*t*_ based on the current belief state *b*_*t*_ to maximize expected payoff:

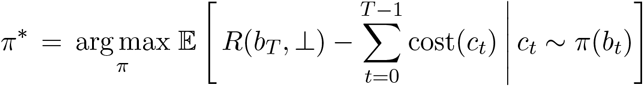

This joint optimization over both when to stop and what to sample can be expressed via the value of computation (VOC). As in (15), we define VOC as the expected increase in total metalevel reward from executing one additional computation *c* and then continuing optimally, rather than terminating immediately:

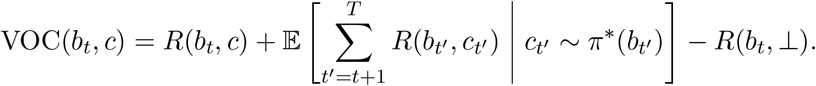

The optimal policy is then defined as selecting computations which have maximal VOC:

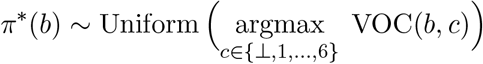

and the optimal policy terminates when no computation has positive VOC.

### Approximating the optimal metalevel policy with Meta-RL

The optimal metalevel policy can be computed analytically using dynamic programming for small belief spaces (16, 64), but these methods are infeasible in larger dimensional spaces such as the present task (in which the belief space has twelve continuous dimensions for an episode’s reward, and six for its relevance). Thus, other methods are necessary for identifying the optimal policy, such as by approximating it using a linear combination of VOC features and fitting to human behavior (15), or by training deep-RL agents on the metalevel MDP (65). Here, we trained a recurrent neural network (RNN) via meta-reinforcement learning to approximate the optimal metalevel policy (40).

The network was implemented as a fully connected gated recurrent unit (GRU) (66) with an actor-critic architecture. The network maintains a hidden state *h*_*t*_ that evolves according to *h*_*t*_ = *f*_*θ*_(*o*_*t*_, *h*_*t−*1_), where *o*_*t*_ is the observation at time step *t* and *θ* denotes the network parameters. The hidden layer consisted of 100 units. The hidden state was provided to two linear output heads. The first was a policy head that produces action logits which are passed through a softmax to define a policy *π*_*θ*_(*a*_*t*_ |*h*_*t*_) over valid actions. Actions are sampled from this policy. The second was a value head which outputs the value baseline *V*_*θ*_(*h*_*t*_).

The agent’s action space consisted of 8 discrete actions: 6 fixation actions (one per episode encoding location on the circle) and 2 decision actions (take or leave). Each time step of the network corresponded to one step of the metalevel MDP. Concretely, the network interacted with the metalevel MDP environment as follows. At each time step *t*, the network received an observation *o*_*t*_, updated its hidden state *h*_*t*_ = *f*_*θ*_(*o*_*t*_, *h*_*t−*1_), and sampled an action *a*_*t*_ ∼*π*_*θ*_(·|*h*_*t*_). If the sampled action was a fixation action *a*_*t*_ = *c* ∈ {1, …, 6}, an episodic memory store (implemented as part of the metalevel MDP environment) returned two pieces of information: (i) a noisy reward sample 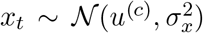 centered on the true reward of episode *c*, and (ii) the episode’s true relevance indicator *s*^(*c*)^ 0, 1 (since *k* = 1; see above). These observations, along with the metalevel reward *r*_*t*_ = *R*(*b*_*t*_, *c*) as defined above, were incorporated into the next time step’s input. If the sampled action was a decision action (take or leave), the trial terminated and the agent received the metalevel termination reward *R*(*b*_*t*_, ⊥). Thus, the network learned to jointly decide which episode to sample and when to stop, with the metalevel MDP’s cost structure shaping the policy through the reward signal.

At each time step, the observation *o*_*t*_ provided to the network consisted of a one-hot encoding of the identity of the fixated episode, the reward sample, the relevance indicator, and the elapsed time since trial onset. Following standard meta-RL practice (37–39), the previous action *a*_*t−*1_ and previous reward *r*_*t−*1_ were also included as inputs, allowing the network’s recurrent dynamics to internalize task structure over the course of a trial. At the start of each trial, the hidden state was initialized to zeros, the fixation pointer was set to blank (no episode attended, represented by an all-zero one-hot vector), the reward sample was set to 0, and the relevance indicator was set to 0.5.

Each training trial corresponded to a single decision in the metalevel MDP. The network was trained using REINFORCE with a baseline to maximize the expected cumulative reward (67). The total loss was a weighted sum of the policy loss *L*_*π*_, the value loss *L*_*v*_, and an entropy regularization loss *L*_*e*_:

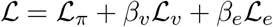

The policy loss used the value baseline to reduce variance:

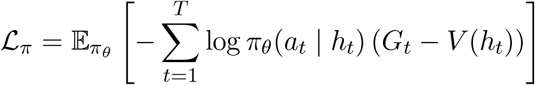

where 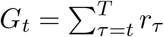 is the empirical return from time step *t* (i.e., the sum of all remaining metalevel rewards in the trial). The value loss trained the baseline to predict the empirical return:

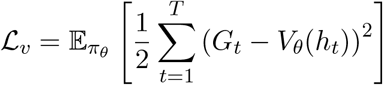

The entropy loss encouraged exploration by penalizing low-entropy policies:

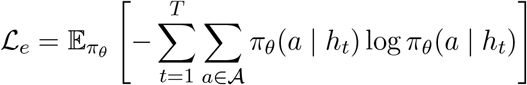

where *A* denotes the set of actions at each time step. The hyperparameters *β*_*v*_ and *β*_*e*_ control the relative contributions of the value loss and entropy regularization, respectively.

We trained the agent on 2.5 × 10^8^ trials. On each trial, the six items were independently sampled from the 16 possible items (defined by four binary features), their rewards were sampled as integers from [− 9, 9] excluding 0, and one of the feature dimensions was selected as the offer. Trials were terminated after a maximum of 100 steps. We used the ADAM optimizer (68) with a learning rate of 10^−3^ and a batch size of 40. To enhance training stability, gradients exceeding 1.0 in magnitude were clipped to 1.0, and all rewards were scaled by a factor of 0.2, which did not affect the optimal policy. The metalevel MDP parameters were set to *σ*_*x*_ = 7, *γ*_sample_ = 0.008, and *γ*_distance_ = 0.04. The network hyperparameters, which control the relative contributions of the value loss and entropy regularization, were set to *β*_*v*_ = 0.05 and *β*_*e*_ = 0.05.

The value of *σ*_*x*_ was informed by participants’ reward recall errors, which had a standard deviation of 4.11. Because the model treats each fixation as drawing a single noisy sample of reward from memory, and a recall report reflects a posterior estimate aggregated over multiple such samples, the per-sample noise *σ*_*x*_ should exceed the observed reward recall error. This is consistent with any sequential sampling account in which averaging over samples yields a more precise estimate than any individual sample. The remaining metalevel MDP parameters (*γ*_sample_, *γ*_distance_) and training hyperparameters (*β*_*v*_, *β*_*e*_) were selected to produce stable training and qualitatively reasonable behavior; these parameters do not have direct behavioral analogues.

#### Network variants

We evaluated two variants of the trained network. In the *no prior memory* variant, the network began each episode with no information about any of the six episodes; it had to fixate to each location to acquire observations about the corresponding episode’s reward and relevance. In the *prior memory* variant, the network received a single noisy reward sample and relevance observation for individual episodes prior to the onset of deliberation. Concretely, we expanded the network’s observation space with 2*N* additional input channels, with *N* for prior reward and *N* for prior relevance. At the start of each trial, *N* episodes were randomly sampled as having prior memory. For each such episode, a reward sample and relevance observation were provided through the expanded input channels. All other reward channels were set to 0, and all other relevance channels were set to 0.5. These additional channels were only used to provide prior memory and their input was set to 0 at subsequent time steps. This variant was designed to model participants entering the decision phase with pre-existing memories for a subset of recalled episodes, consistent with the offer cue triggering associated retrieval before deliberate sampling begins. We set *N* = 5 to approximate the number of episodes likely to be accessible by participants at the onset of deliberation, as participants recalled an average of 4.62 ±0.14 items per round during the post-decision free recall task.

For each variant, we trained 5 networks with different random seeds. During evaluation, network parameters were frozen and all adaptation occurred through the recurrent dynamics alone (38). Each network was evaluated on independently sampled episodes, and results were aggregated across the five seeds.

### Evidence Accumulation Analysis

To examine how the network accumulates evidence during deliberation, we computed two decision variables at each time step. For the network, we defined the decision variable as the difference between the take and leave output logits (DV_*t*_ = logit_take,*t*_ −logit_leave,*t*_), which reflects the network’s log-odds of taking versus leaving prior to committing to a decision. Because each trained network seed has a different initial logit bias, we baseline-subtracted each trial’s decision variable by its value at *t* = 0 (before any fixation-specific information has been processed), so that all seeds begin at zero. For the metalevel MDP, the decision variable was defined as the Bayesian offer value estimate, *V*_*t*_ = ∑_*i*_ *µ*_*t,i*_*· ρ*_*t,i*_, where *µ*_*t,i*_ is the posterior mean reward belief at location *i* and *ρ*_*t,i*_ is the posterior probability that location *i* is relevant to the current offer. Both decision variables were computed at each fixation step and averaged separately for take and leave trials. Fixation steps were binned as 0 through 6 individually and 7+, with the latter bin aggregating all fixation steps from 7 onward. This analysis was performed on the *no prior memory* network, in which the network begins each trial with no information about the episodes and must discover their rewards and relevance entirely through deliberate sampling.

### Belief State Decoding

To determine whether the network’s hidden state encodes the sufficient statistics of the metalevel MDP’s Bayesian belief state at each timepoint, we trained linear decoders to predict each belief component from the network’s 100-dimensional GRU hidden state. We decoded four target variables: the offer value estimate *V*_*t*_ and the three per-location sufficient statistics of the belief (the posterior mean reward *µ*_*t,i*_, the posterior precision *λ*_*t,i*_, and the posterior relevance probability *ρ*_*t,i*_) for each of the six locations. Each decoder was a linear regression trained to predict each variable from the 100-dimensional hidden state. Decoding performance was assessed using 5-fold cross-validated *R*^2^ under a group-aware split, with each trial assigned entirely to a single fold so that timesteps from a given trial never appeared in both train and test. For the per-location variables (*µ*_*t*_, *λ*_*t*_, *ρ*_*t*_), we report the mean *R*^2^ across the six locations. Chance performance was estimated by permuting the target variable and re-fitting the decoder. All analyses were conducted independently for each of five network seeds.

### Comparison of network behavior and gaze with humans

#### Overall performance

We conducted several analyses in order to determine whether humans and the network exhibited similar patterns of behavior and fixations. For each comparison analysis, we averaged across the five random seeds fit per agent to obtain a single reference value, subtracted this value from each human participant’s value, and asked whether the resulting deltas differed from zero. All regression models were fit using methods identical to prior analyses.

We first compared overall performance on the decision making phase by computing the network’s accuracy *a*_nn_ as the mean across seeds. We then subtracted this measure per person (*δ*_acc,*i*_ = *a*_*i*_*−a*_nn_) and fit a simple linear regression to assess differences from zero: *δ*_acc,*i*_ = *β*_0_ + *ε*_*i*_. We followed an identical approach to assess differences in the proportion of fixation time to relevant items as well as in the proportion of fixation time directed to relevant items during either initial fixations to item locations (up to 6) and revisits (any fixation occurring after the first).

#### Valence and decision interactions

To examine whether the network displayed any differences from people in the interaction between the proportion of time spent fixating on item locations as a function of their valence and choice type, for each participant we computed:

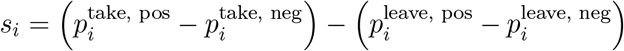

where 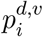 denotes the mean proportion of fixation time to relevant items with valence *v* under decision *d* for subject *i*. We then computed this same score for the network and followed an identical approach to assess differences from human participants.

#### Fixation counts

We also compared the total number of fixations per trial between humans and the network. For each participant, we computed the mean number of fixations per trial to offer-relevant and offer-irrelevant locations separately, subtracted the corresponding network benchmarks, and fit a mixed-effects model predicting these deltas from a contrast-coded relevance indicator (*R*_*i*_) with a random intercept per participant:

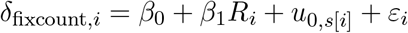

The intercept, scaled by 2 to recover the total difference in fixation count per trial, captures the overall difference between humans and the network, and the slope captures whether this difference is disproportionately larger for offer-relevant versus offer-irrelevant items.

#### Fixation dynamics over deliberation

We additionally conducted several analyses comparing the evolution of fixation durations as well as where fixations occurred over the course of deliberation. We first aimed to assess where there were differences between humans and the network in both 1) the distribution of fixation durations and 2) the number of fixations that accounted for 80% of cumulative fixation time. To accomplish these goals, we calculated the area under the cumulative fixation curve (AUC) as well as the ordinal fixation position as which cumulative fixation time first reached 80%. We calculated these scores for both each participant and the network, and assessed their difference using an identical approach to the above.

We then asked how fixation allocations evolved over the course of deliberation by computing two position-specific measures: i) the mean proportion of fixations to relevant items at position *j* for participant *i*, and ii) the proportion of fixation time directed to positively and negatively valenced relevant items at each fixation position *j* for each participant *i*. For each measure, we subtracted the corresponding network benchmark (averaged across seeds at each position) and fit a mixed-effects regression model.

For the proportion of relevant fixations, we fit:

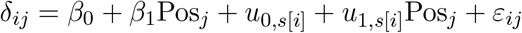

For the analysis of valence-dependent fixation proportions at each position, both decision type (Dec; take = +0.5, leave = −0.5) and valence were contrast-coded (Val; positive = +0.5, negative = −0.5). We then fit a model including all two- and three-way interactions with position:

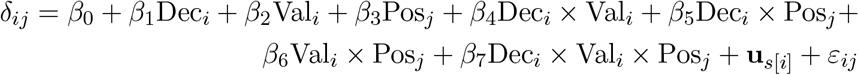

where **u**_*s*[*i*]_ includes random intercepts and random slopes for decision, valence, and position per participant.

### Choice prediction comparison

We next compared the network’s cross-validated choice prediction accuracy from the *L*_2_-regularized logistic regression (described above) to that of human participants. The same model was fit to the network’s simulated fixation data, yielding a single CV accuracy estimate. We assessed whether this value fell within the 95% confidence interval of the human CV accuracy. We performed this comparison twice: once using the network’s full fixation data, and once after randomly dropping a proportion of the network’s fixations calibrated to the mean proportion of fixation time that humans allocated to item locations at significant free recall time-points, in order to simulate the imperfect correspondence between fixations and memory sampling. Note that to ensure the correspondence in CV accuracy between humans and the network after dropping fixations was specific to the proportion measured by the free recall task, we repeated this analysis several times, maintaining a different proportion of fixations on each iteration (**Fig. S4**).

#### Evaluating how fixations were generated

To characterize the factors that drive which item location participants and the network fixated next (relative to the current location upon which they were fixating), we modeled each next-fixation event as a categorical choice over the set of candidate locations *C* that could be fixated next, defined as all locations that were fixated at any point during each trial. For each fixation at which the currently fixated location was *i*, we treated the next fixated location *j* as a draw from a softmax over the remaining candidate locations:

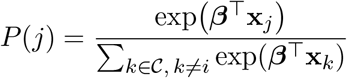

where **x**_*k*_ is the vector of predictors for candidate *k* and ***β*** are the coefficients to be estimated. Trials for which fewer than two unique items were fixated were excluded from this analysis. We modeled this as a conditional logistic regression with ten predictors that varied per-location on each trial, organized into three categories. Each predictor was *z*-scored prior to model fitting to allow coefficients to be compared on the same scale.

##### Resource-rational predictors

We first aimed to assess whether predictors aligned with a resource-rational policy influenced where each fixation landed. We identified three such predictors. The first, which was the *inverse relevant fixation time*, was defined as 1*/*(1 + *t*_*k*_) for offer-relevant candidate locations, where *t*_*k*_ is the cumulative time spent fixating location *k* up to that point in the trial, normalized by the dataset-wide mean cumulative fixation time, and the 1 in the denominator keeps the predictor finite for locations not yet fixated. We normalized in this way because cumulative fixation time is recorded in milliseconds for participants but in discrete steps for the network and our null models, so dividing by the dataset mean places each predictor on a common scale across fits. This predictor is largest for offer-relevant locations that have not yet been fixated and falls toward zero as fixation time at a location accumulates. It is therefore proportional to uncertainty reduction in the metalevel-MDP model, as more time spent fixating indicates more time sampling. This reduction is valuable only for offer-relevant episodes, whose rewards alone determine the offer’s value. Because the model predicts that an episode’s reward uncertainty decreases the more it is sampled, an agent following a resource-rational policy should yield a positive coefficient for this predictor.

The second and third predictors were the *clockwise* and *counter-clockwise distances* between the currently fixated location and each candidate. Because the six episode locations were arranged around a circle, every candidate could be reached by moving either clockwise or counter-clockwise from the current location. The clockwise distance was the number of steps to a candidate in the clockwise direction when it lay no more than three steps that way, and zero otherwise; the counter-clockwise distance was defined symmetrically. Because a resource-rational policy incurs a cost proportional to the distance moved between successively sampled locations, an agent following such a policy should favor candidates near the current location, yielding negative coefficients for both predictors.

##### Encoding-order predictors

We next aimed to assess whether the order in which episodes were studied influenced fixation choice, which is not captured by the resource-rational policy. We included four such predictors. The *forward* and *backward temporal distances* were, respectively, how many positions later or earlier in the study sequence a candidate had been encoded relative to the currently fixated location. *Primacy* and *recency* were binary indicators of whether a candidate was the first or last episode encoded on that trial. Because the resource-rational policy is indifferent to study order, it predicts no effect of these predictors, and a non-zero coefficient on any of them would instead reflect an influence on fixation arising from how memory is organized by temporal properties at encoding time.

##### Biases

Finally, we aimed to assess whether fixation choice was pulled by factors with no clear role in episodic sampling. We included three such predictors. The *reward magnitude* and *signed reward* predictors were the absolute and signed reward values of a candidate episode. Under the resource-rational policy these should carry no weight, because the value of sampling a location depends on the reward uncertainty it would resolve, not on the magnitude or sign of the reward itself (see **supplementary text**). The *previous item* predictor was a binary indicator of whether a candidate was the location fixated immediately before the current one (a return to the just-fixated location), which reflects “alternating” behavior between two locations; its interpretation is discussed separately in the **supplementary text**. Non-zero coefficients on these predictors indicate influence of biases on fixations beyond encoding order and resource-rational sampling.

For human participants we fit a mixed effects version of this model with participant-level random slopes on all ten predictors. We placed a half-normal prior ***τ***∼*N* ^+^(0, 1) on the random-slope standard deviations, an LKJ(2) prior on their correlation matrix, and weakly-informative *N*(**0**, 1.5^2^) priors on the population-level coefficients ***β***. For the network and the null models we fit fixed-effect-only versions of the same model, omitting the random-slope terms. Model fitting was performed using No-U-Turn Sampling (60) in Stan. Iterations and convergence criteria were identical to all other Bayesian regression models.

##### Null models

We ran the above analysis on two different null models to ensure that effects were not due to properties of the task or trivial behavior. The first was a *uniform-random* model which drew each fixation uniformly at random from the locations other than the current one. The second was an *adjacent* model which instead stepped to one of the two locations neighboring the current one on 90% of fixations, and to a randomly chosen location on the remaining 10% of fixations, producing sequences with spatial structure but no uncertainty-driven or memory-based component. Both null models were matched to the number of fixations participants made on each trial and resampled ten times per trial.

#### Analyses of spatial transition structure in fixations

Finally, we conducted two sets of analyses to further assess spatial influences on next fixation generation. First, we asked whether participants showed any spatial biases in their fixation patterns. Specifically, we computed a 6 × 6 row-normalized transition matrix from each participant’s fixation position sequences. We compared each observed transition matrix to three idealized sweep templates defined on the circle: *Bidirectional* from each position *i*, transition to (*i* + 1) mod 6 or (*i* − 1) mod 6 with equal probability; *Forward* from each position *i*, always transition to (*i* + 1) mod 6; and *Backward* from each position *i*, always transition to (*i* − 1) mod 6.

We quantified similarity between each of these template matrices and the observed transition matrix as the Pearson correlation between them. To establish a chance baseline, we computed a within-trial shuffled null distribution (1000 permutations) that preserved the set of visited positions per trial, but shuffled their temporal order. We computed the similarity between each of the 1000 permutations and defined *δ*_similarity_ as the observed similarity minus the mean shuffle similarity. We then fit the following mixed effects model:

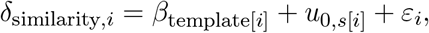

where *β*_template[*i*]_ is a fixed effect for each template direction (bidirectional, forward, backward), and the intercept is suppressed so that each coefficient directly estimates the mean *δ*_similarity_ for its respective template. We examined whether each template’s coefficient differed from zero (i.e., whether observed fixation transitions resembled that sweep pattern more than expected by chance) and then conducted pairwise contrasts between all three template directions.

Second, we characterized the extent to which fixations followed consecutive *sequences*, which we defined as transitions to adjacent positions in the same rotational direction (e.g., positions 2 → 3 → 4 constitutes a sequence of length 3). For each trial, we computed the fraction of all non-zero transitions that belonged to sequences of each length (binned as lengths 1, 2, 3, 4, 5, and ≥6). The chance-corrected delta was defined as the observed fraction minus the mean fraction under the shuffled null distribution. We then fit the following mixed effects model:

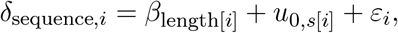

where *δ*_sequence_ is the chance-corrected fraction of transitions belonging to sequences of a given length for observation *i*, and *β*_length[*i*]_ is a fixed effect for each sequence length bin (1, 2, 3, 4, 5, ≥6), with the intercept suppressed so that each coefficient directly estimates the mean *δ*_sequence_ for its respective bin. We again examined whether each bin’s coefficient differed from zero, which would indicate that sequences of that length occurred more (or less) frequently than expected by chance.

## Acknowledgments

The authors thank Natalie Biderman, Fred Callaway, Camilla van Geen, and members of the Mattar Lab for insightful conversations and comments on the manuscript. This work was funded by the National Science Foundation SBE Postdoctoral Research Fellowship grant # 2507527.

## Author Contributions

Conceptualization: JN, MM; Methodology: JN, SC, MM; Formal Analysis: JN, SC; Investigation: JN; Visualization: JN; Software: JN, SC; Funding acquisition: JN; Project administration: JN; Supervision: MM; Writing – original draft: JN; Writing – review & editing: JN, SC, MM

## Data, code, and materials availability

All data, code, and materials can be found on [github](https://github.com/jonathanicholas/ncm2026_optimalepisodic).

## Supplementary Text

### Reward magnitude does not impact the optimal sampling policy

Neither humans (*β*_|*r*|_ = +0.019, 95% HDI [−0.018, 0.054]; *β*_*r*_ = +0.015, 95% HDI [−0.015, 0.047]) nor the trained network (*β*_|*r*|_ = − 0.019, 95% HDI [− 0.022, − 0.016]; *β*_*r*_ ≈ 0, 95% HDI [− 0.003, 0.002]) showed a substantive positive effect of reward magnitude or signed reward on which items were fixated in our conditional logistic regression analysis. This may seem surprising on two counts. First, it is intuitive to think that a sample of a high-magnitude item is more decision-relevant than a sample of a low-magnitude item because the former contributes more to the offer’s value. Second, in a closely related framework applied to multi-alternative choice, Callaway et al. (15) showed that it is optimal to preferentially sample items that are currently estimated to be highest in value.

Counter to this intuition, magnitude-blindness is a structural prediction of our metalevel MDP that arises from the form of the termination reward. Here we provide an overall argument and derivation for why this is the case.

First, the value of sampling an item is governed by *how much the sample could change an agent’s decision*, not by how rewarding the item is currently believed to be. In our task, the decision is take-or-leave based on whether the relevance-weighted sum of item rewards is positive. An agent’s current best guess for that sum already incorporates every item’s currently-believed reward. Sampling cannot *add* an item’s magnitude to the decision, because that magnitude is already baked in. What sampling can do is *sharpen* an agent’s belief, and the size of the typical sharpening per sample is determined by how uncertain the agent currently is.

To make this concrete, suppose item A has prior *N*(*µ* = 1, *σ* = 1) and item B has prior *N*(*µ* = 7, *σ* = 1). After one sample, item A’s posterior mean lands within roughly ± 0.7 of 1 and item B’s lands within roughly ± 0.7 of 7, making the spread of the potential shift identical. The change in the decision variable produced by one sample is therefore identical regardless of which item is sampled, and the optimal sampling policy cannot prefer one item over the other on the basis of magnitude alone. We next provide a derivation to demonstrate this point formally.

We use the same notation introduced in **Materials and Methods** and quickly review it here for completeness. The belief state *b*_*t*_ = (***µ***_*t*_, ***λ***_*t*_, ***ρ***_*t*_, last_*t*_, *f*) represents Gaussian reward posteriors and offer-relevance probabilities for each of the *N* = 6 episodes. The decision variable summarizing the agent’s current valuation of the offer is

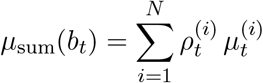

and the metalevel termination reward is *R*(*b*_*t*_, ⊥) = max {*µ*_sum_(*b*_*t*_),0} . We assume (consistent with our implementation and the empirical justification in **Fig. 4**), that relevance is known and takes values in {0, 1} .

We focus here on establishing that magnitude does not enter the optimal policy under a one-step look-ahead approximation of the value of computation (VOC), which is tractable in closed form, but the same conclusion should extend to the full multi-step VOC.

The one-step VOC for sampling episode *c* at belief state *b*_*t*_ is the expected gain in termination reward from one additional sample,

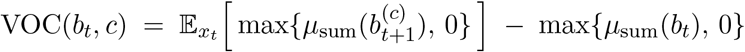

A Gaussian update with prior *N* (*µ*_*c*_, 1*/λ*_*c*_) on episode *c*’s reward and observation 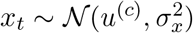 yields the posterior reward mean

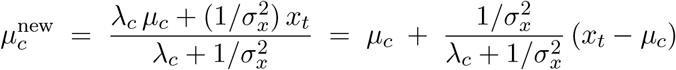

Sampling *c* shifts only *µ*_*c*_, so the change in the decision variable from sampling *c* is

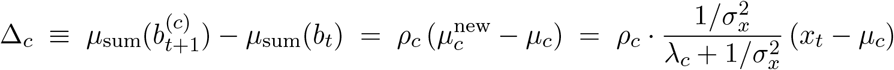

An agent must evaluate VOC for each candidate before observing *x*_*t*_, which requires the predictive distribution of Δ_*c*_. This is fully characterized by the mean and variance, which are obtained by marginalizing the current belief about the true reward *u*^(*c*)^ ∼ *N* (*µ*_*c*_, 1*/λ*_*c*_) over the observation likelihood 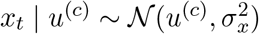

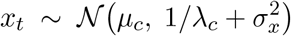

whose mean is the agent’s current best guess for the reward and whose variance combines belief uncertainty and observation noise. Under this predictive, E[*x*_*t*_ − *µ*_*c*_] = 0, so Δ_*c*_ has expected value zero. Its variance is

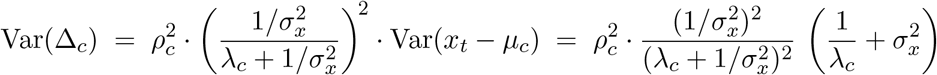

which simplifies to

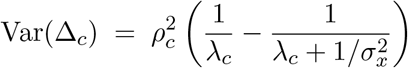

The variance depends only on *ρ*_*c*_, *λ*_*c*_, and 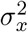, but notably not on *µ*_*c*_. Substituting into VOC,

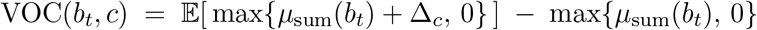

which depends on the belief state through only two quantities: *µ*_sum_(*b*_*t*_), which is identical for every candidate episode at a given timepoint, and Var(Δ_*c*_). Two candidates with identical *ρ*_*c*_ and *λ*_*c*_ therefore have identical one-step VOC, regardless of their reward magnitudes.

Because the metalevel MDP’s terminal reward, transition dynamics, and costs all depend on the individual reward means 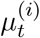 only through their relevance-weighted sum *µ*_sum_(*b*_*t*_), the optimal policy depends on ***µ***_*t*_ only through *µ*_sum_. Two candidates with identical relevance and precision should therefore have identical multi-step VOC at any belief state, regardless of their individual reward beliefs, allowing this conclusion to extend to the full multistep VOC defined in the **Materials and Methods**.

### Contrast with Callaway et al., 2021

The difference in the effect of magnitude compared with (15) is due to the form of the termination reward used in either setting. In (15), the agent makes a multi-alternative choice and the termination reward is 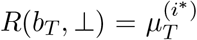 where *i*^*^ = arg max_*i*_ 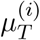, or the value of the chosen item. The argmax operator makes the termination reward *non-linear* in ***µ***, so a sample’s expected value depends on whether it can plausibly change which item is the argmax. Items far below the best item cannot become it and have near-zero VOC; items near the best can flip the choice and have higher VOC. The agent’s preference for sampling high-*µ* items is the consequence of this rule.

In our setting, the termination reward is instead max{*µ*_sum_, 0}, a *linear* combination of item-level beliefs (clipped at zero). A constant shift in any *µ*_*c*_ produces an equal shift in *µ*_sum_, but the choice between candidate episodes depends only on how much the spread of possible *µ*_sum_ can be reduced by sampling each one, which is (as shown above) not a function of reward magnitude.

### Previous-item effects reflect differential spatial and uncertainty sensitivities

Our conditional logistic regression also included a *previous item* predictor (a binary indicator of whether a candidate location was the one fixated immediately before the current location) as a candidate measure of return tendency (perseveration or rehearsal). Participants and the network showed different effects on this predictor. Participants had a positive coefficient (*β*_human_ = +0.088, 95% HDI [0.027, 0.144]; **Table S7**), suggesting that they often preferred to return to the just-sampled location, whereas the network had a negative coefficient (*β*_network_ = − 0.127, 95% HDI [− 0.130, − 0.125]; **Table S7**), suggesting that it actively avoided its preivous location.

These effects are not included in the main text because they are likely byproducts of the two resource-rational factors operating to different degrees in people and the network: sensitivity to spatial structure and sensitivity to reward uncertainty. Spatial structure alone is sufficient to produce a positive previous-item coefficient. Specifically, an adjacent null model, which transitions only between neighboring locations on the circle (**Fig. S5**; **Table S7**), itself yields a coefficient of comparable magnitude to participants’ (*β* = +0.075). This follows from the geometry of the task: stepping between adjacent locations on a six-location ring inevitably returns to the previous location more often than chance. Because participants weighted spatial proximity more strongly than the network (**Fig. S6**; **Table S8**), this same spatial structure plausibly accounts for their apparent return tendency.

Likewise, the network’s negative coefficient instead reflects its uncertainty-driven sampling. Because the network prefers whichever location is currently most uncertain, and the just-sampled location is unlikely to be among the most uncertain, the policy avoids it. This uncertainty preference overrides the network’s own spatial preference and produces the observed negative coefficient. The network also showed stronger uncertainty-driven effects than human participants (i.e. the magnitude of its inverse relevant fixation time coefficient was larger than the fixed-effect coefficient for people), explaining why this effect appears to be in the opposite direction from participants.

We therefore interpret these differential effects not as separate biases, but as natural byproducts of joint sensitivity to spatial structure and reward uncertainty: participants’ positive coefficient is dominated by the spatial-structure component, while the network’s negative coefficient is dominated by the uncertainty component.

## Supplementary Figures and Tables

**Figure S1:**
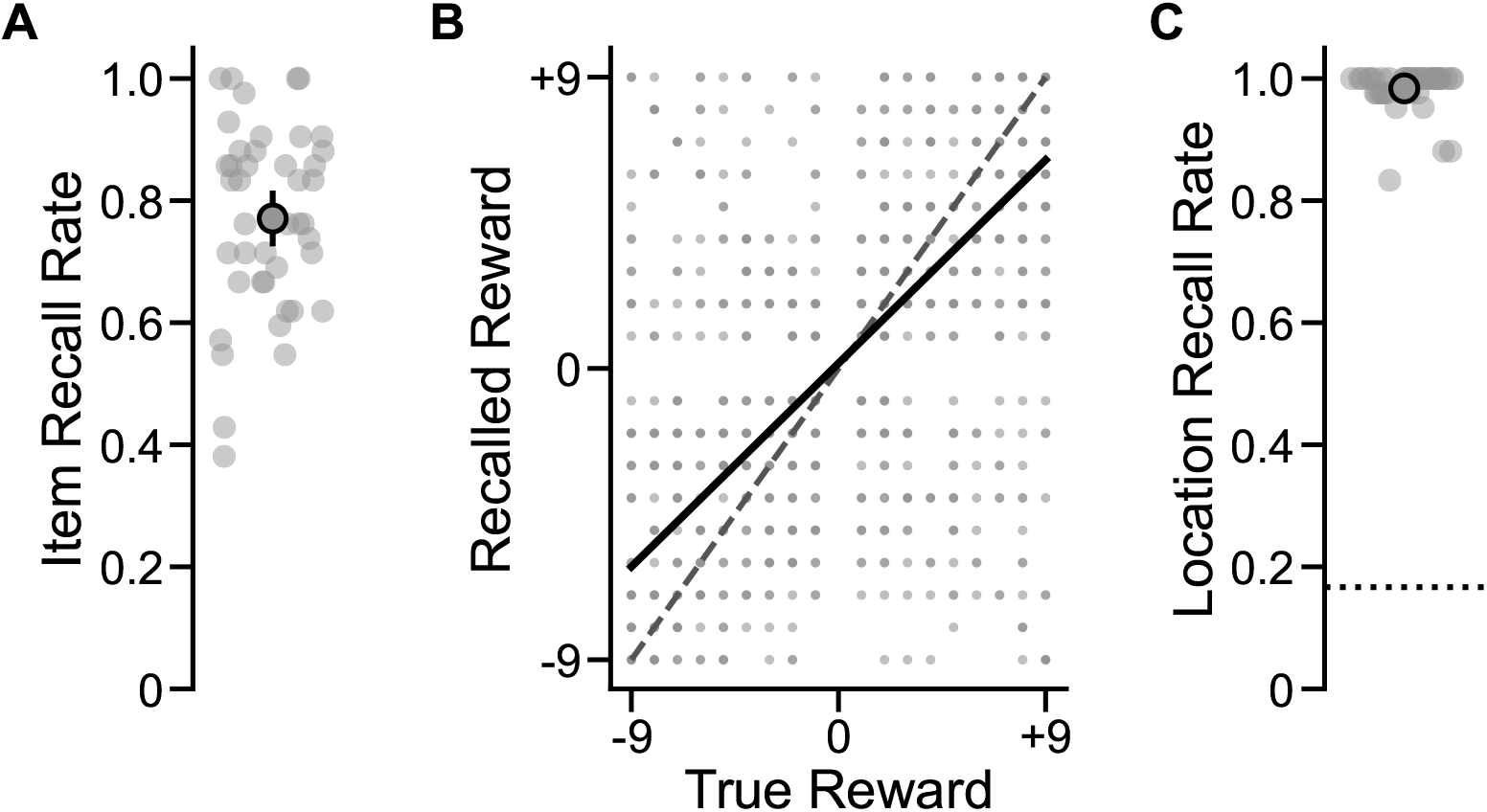
Memory phase performance. (A) Performance on the free recall phase. Item recall rate is the number of items (out of 6) that were correctly recalled on each round. (B) Performance on the reward recall phase shown as the relationship between the true reward associated with each item and participants’ recollection of these rewards. (C) Performance on the location recall phase. Location recall rate is the number of locations (out of 6) that were correctly recalled on each round.

**Figure S2:**
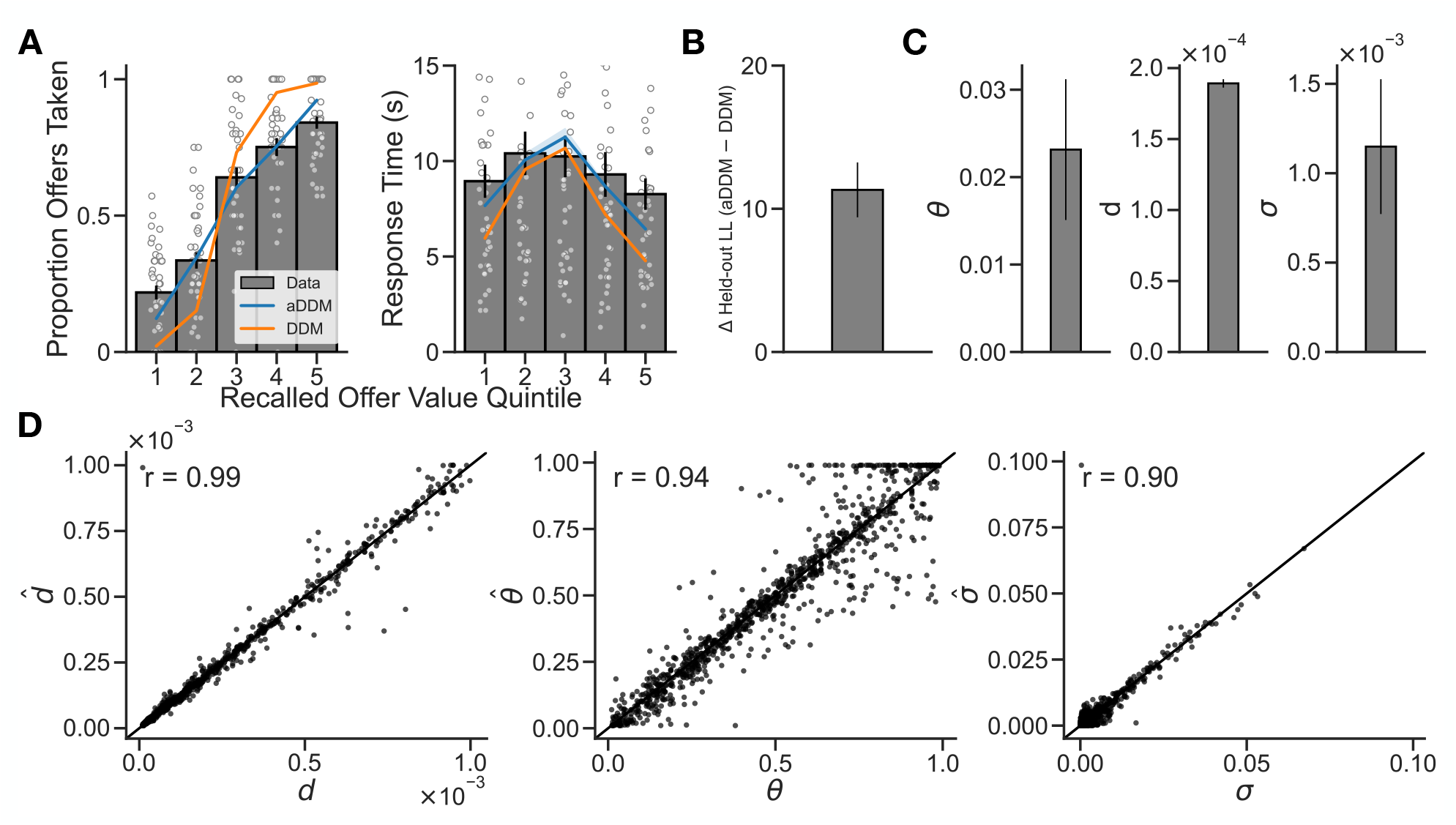
Attentional drift-diffusion model (aDDM) fits, model comparison, and parameter recovery. (A) Posterior predictive checks showing proportion of offers taken (left) and response time (right) as a function of recalled offer value quintile. Gray bars and individual data points show observed data; blue and orange lines show predictions from the aDDM and a standard DDM (aDDM with *θ* fixed to 1), respectively. The aDDM provides a closer estimate in both cases. (B) Difference in held-out log-likelihood between the aDDM and DDM (Δ*loglik*_aDDM−DDM_) from 10-fold cross-validation. Positive values indicate better out-of-sample prediction by the aDDM. Error bars denote standard error. (C) Group-level maximum-likelihood parameter estimates for the aDDM. *θ*: attentional bias parameter controlling the relative weighting of fixated versus non-fixated relevant items (lower values indicate stronger attentional modulation); *d*: drift scaling factor linking attended offer value to the rate of evidence accumulation; *σ*: diffusion noise scale. Error bars denote standard error. (D) Parameter recovery. Each panel plots recovered 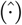 against true parameter values from 1000 simulated datasets spanning the parameter space. Pearson correlations are shown in each panel. The identity line is plotted for reference.

**Figure S3:**
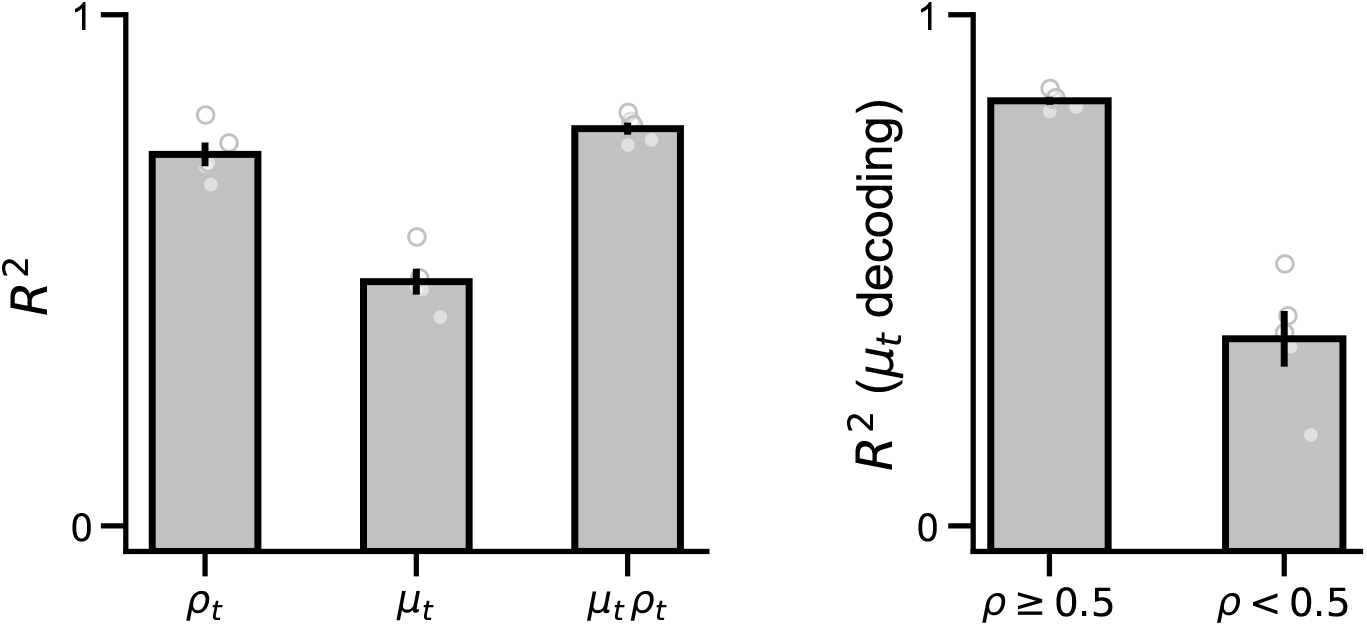
Decoding of belief sufficient statistics from network hidden state reflects their contribution to *V*_*t*_. Linear decodability of each per-location belief component reflects how strongly it contributes to the policy’s value estimate *V*_*t*_ = Σ_*i*_*µ*_*t,i*_*ρ*_*t,i*_. Bars show cross-validated linear decoding *R*^2^ from the network’s hidden state, averaged across the six episode locations; circles show per-seed values. (**Left**) Per-location decoding of the relevance posterior *ρ*_*t,i*_, the posterior mean reward *µ*_*t,i*_, and their product *µ*_*t,i*_*ρ*_*t,i*_, which is the per-location quantity that contributes to *V*_*t*_. (**Right**) Per-location decoding of *µ*_*t,i*_ restricted to timesteps where *ρ*_*t,i*_ ≥0.5 (location likely relevant to the offer) versus *ρ*_*t,i*_ *<* 0.5 (location likely irrelevant). *µ*_*t,i*_ is decoded substantially better in relevant locations, where it contributes to *V*_*t*_. Together, these analyses indicate that the network most cleanly represents the quantities that the policy actually reads out, as it preserves the per-location relevance-weighted reward *µ*_*t,i*_*ρ*_*t,i*_ that contributes to *V*_*t*_, while *µ*_*t,i*_ itself is encoded primarily as part of this product rather than as a separate factor.

**Figure S4:**
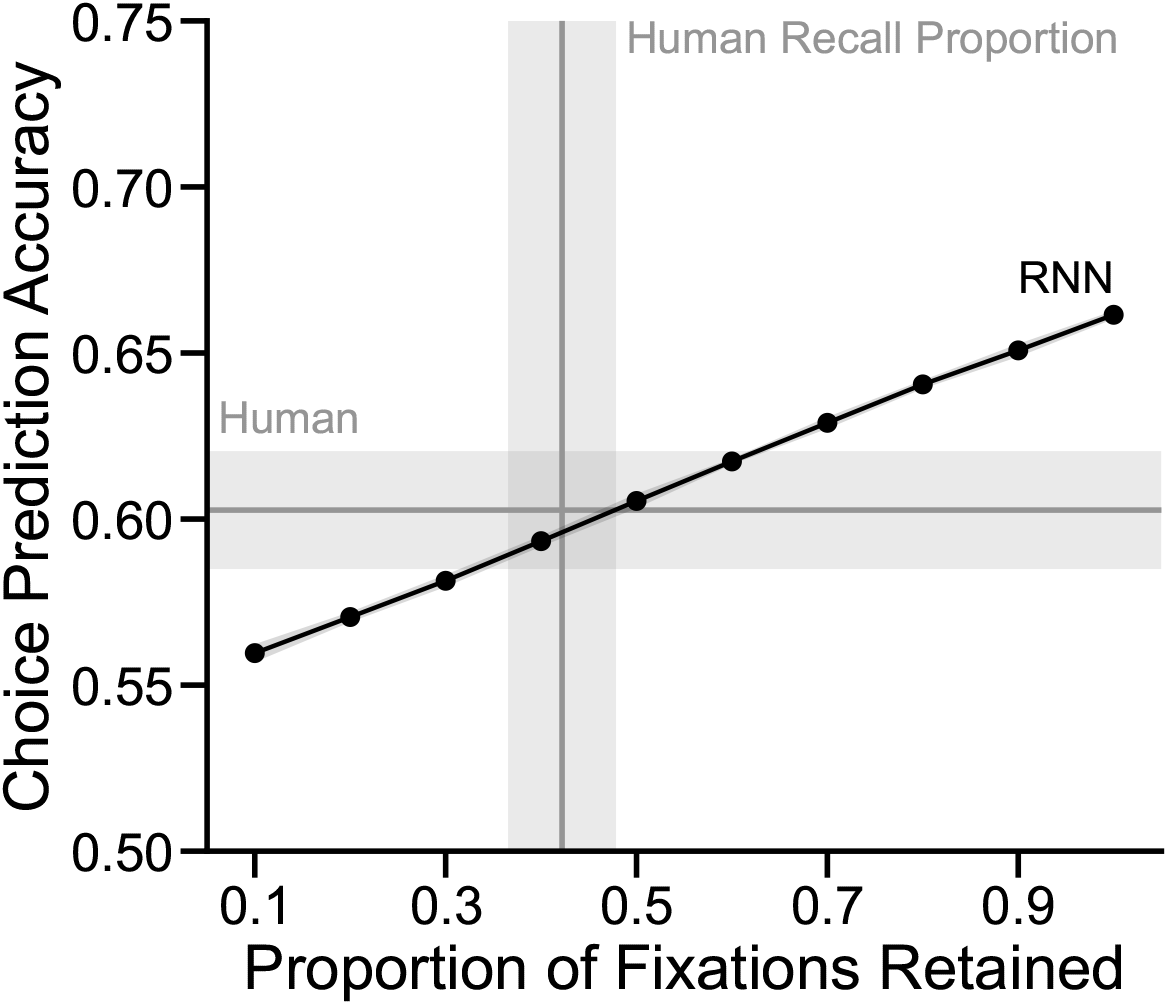
Effect of dropping RNN fixations on choice prediction accuracy. Choice prediction accuracy from a 10-fold cross-validated logistic regression that predicts choices from fixation-derived features (proportion of fixation time at each location, interacted with reward value and relevance) as a function of the proportion of fixations retained. At each retention level, individual fixations were randomly dropped before recomputing features and re-evaluating cross-validated prediction accuracy. The black line and shaded region show mean accuracy ±95% confidence intervals across folds for the network’s fixation data. The horizontal gray line and band show human choice prediction accuracy (±95% confidence intervals) from the same logistic regression applied to the human fixation data. The vertical gray line and band show the mean proportion of fixation time (±95% confidence intervals) that humans devoted to items at timepoints that significantly predicted subsequent recall. As the proportion of fixations retained decreases, prediction accuracy declines monotonically, approaching human-level accuracy near the empirical human recall proportion.

**Figure S5:**
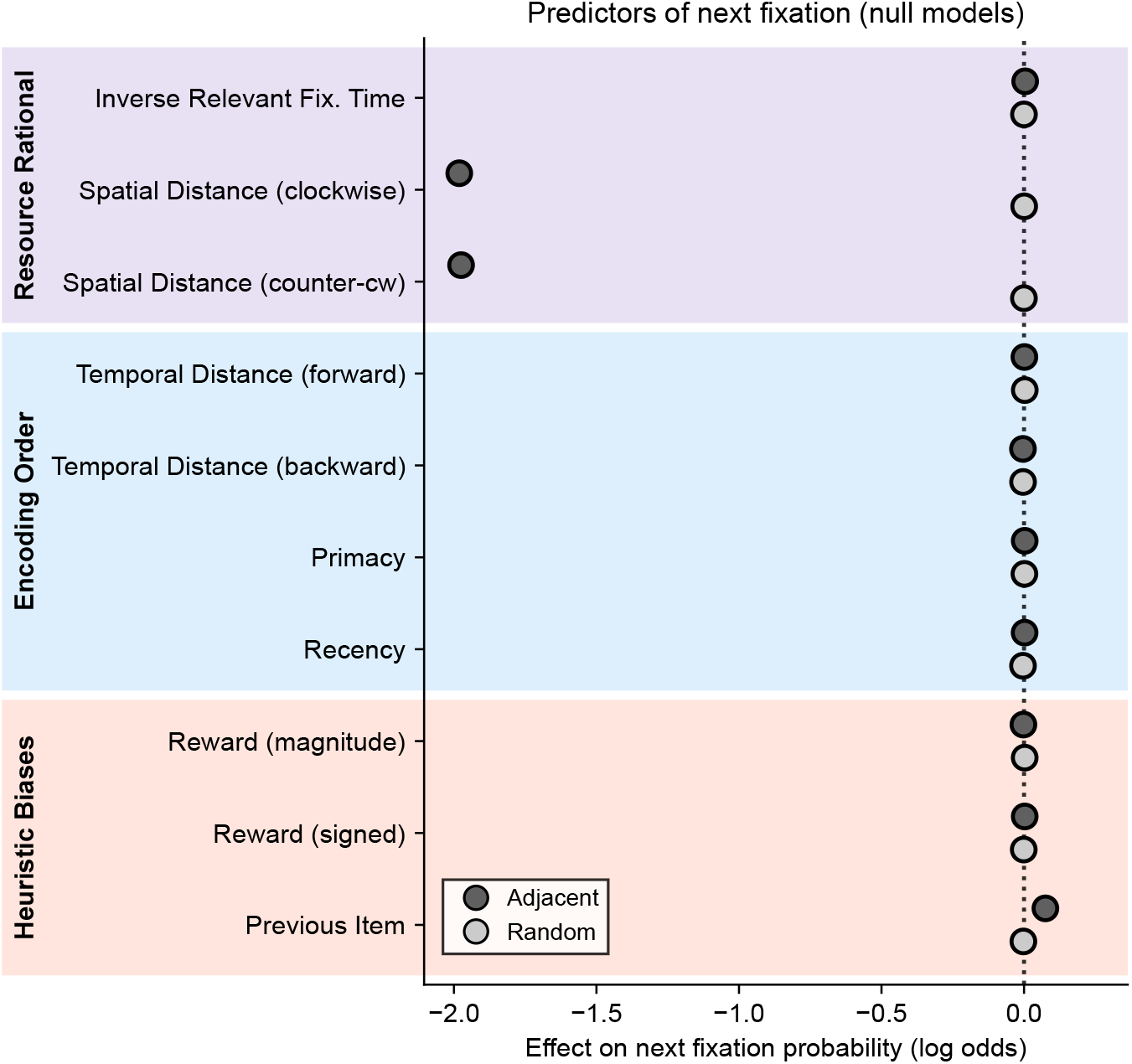
Next-fixation conditional logistic regression fit to null model. The next-fixation model (**Fig. 5**) fit to two null sampling models, each matched to participants’ trial-length distribution and resampled ten times: an adjacent sampler that steps to a neighboring location with occasional random jumps, and a uniform-random sampler. Points and horizontal lines show the fixed-effect posterior mean and 95% HDI for each of the ten *z*-scored predictors, organized into the same three categories as **Fig. 5**. The adjacent null produces large spatial-distance effects and a modest previous item bias, as expected from its construction (see **supplementary text**), but neither null shows the inverse-relevant-fixation-time effect or any encoding-order structure. The dotted line at zero indicates no effect.

**Figure S6:**
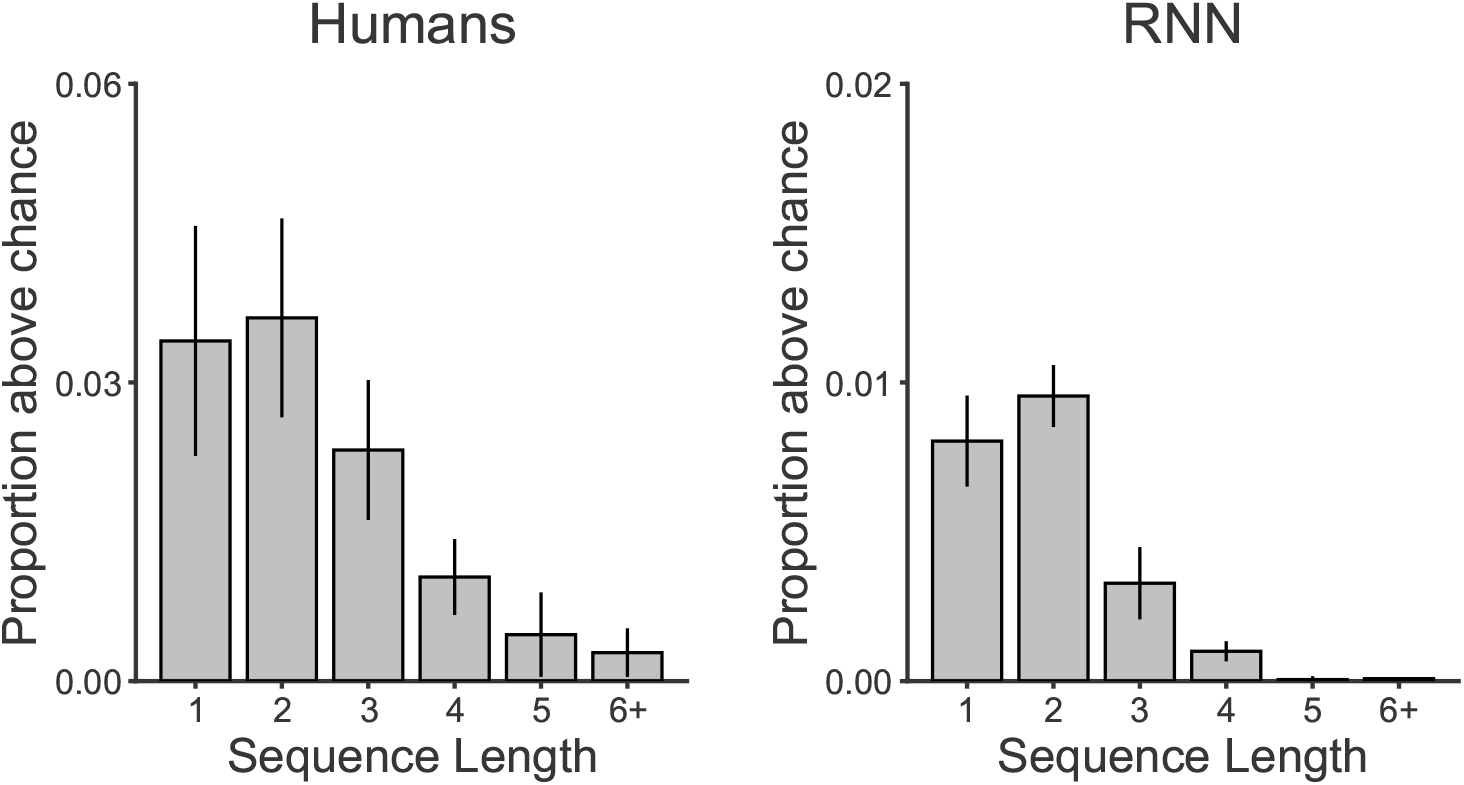
Sequential structure in the order of episodic sampling. Chance-exceeding proportions of sequences of fixations that transitioned between adjacent locations in the same circular direction. Both participants and the RNN moved predominantly between adjacent encoding locations, with successive transitions often continuing in the same circular direction. This structure follows from the optimal sampling policy: a decision maker should remain at an episode only while continued sampling reduces uncertainty about its reward more than would sampling an episode she has spent less time on, and when she switches she should prefer adjacent locations to minimize transition costs. An emergent consequence is that she continues sampling in the same direction around the circle, since the location ahead is typically one she has sampled less and therefore knows less well. The network reproduces these patterns by learning to trade off information gain against transition cost at each switch, indicating that the same uncertainty-reducing policy gives rise to the sequential structure seen in participants. Participants nonetheless showed stronger effects than the network.

**Figure S7:**
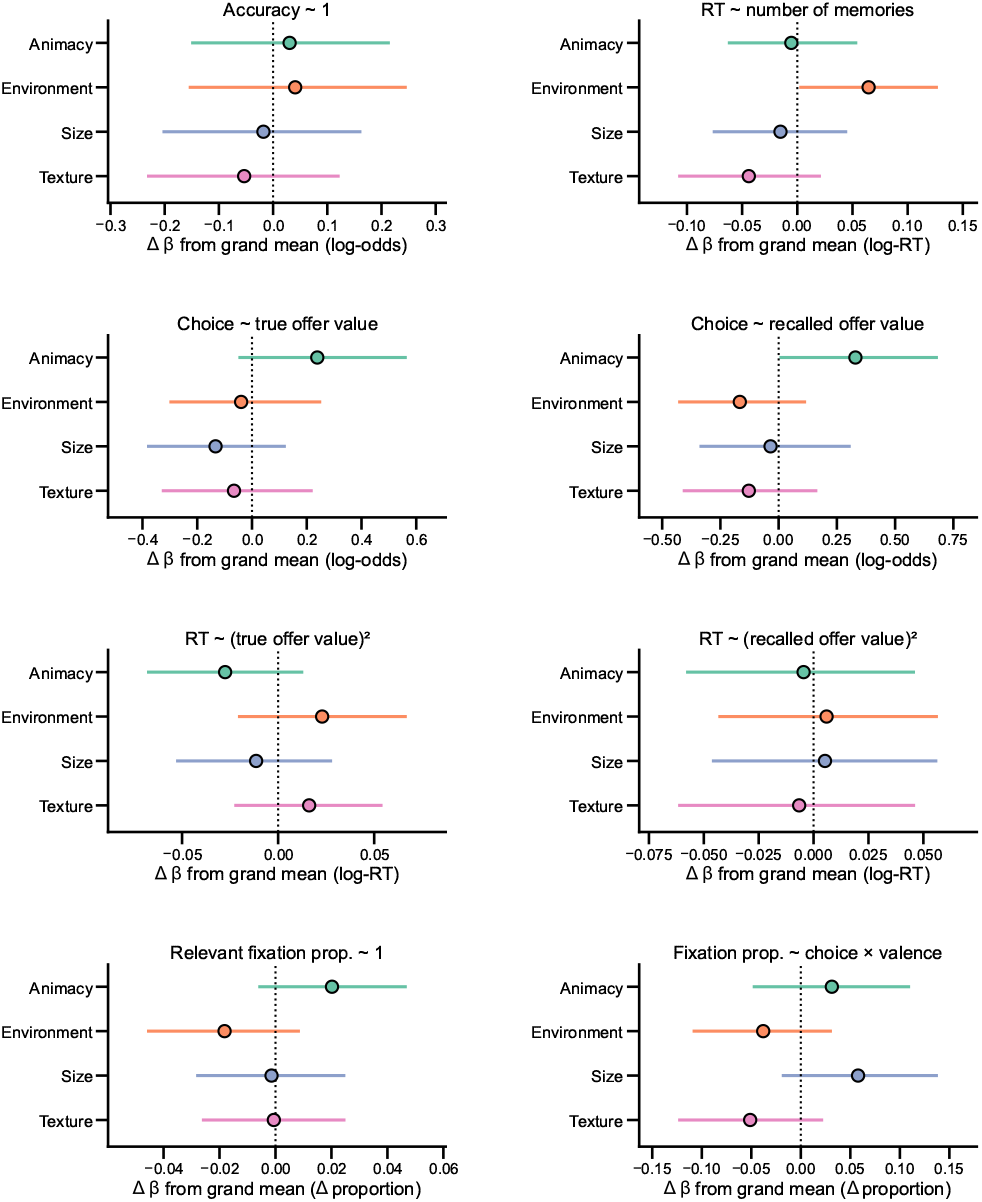
Behavioral and eyetracking results by feature. To assess whether overall effects were driven by a subset of the four features, we reparameterized several analyses so that each feature dimension (animacy, environment, size, texture) received its own coefficient (via an interaction term) that varied by subject, with all other model aspects preserved. These per-feature reparameterizations covered overall choice accuracy (Fig. 1B), the effect of the number of memories on RT (Fig. 1C), the effect of true and recalled offer value on choice (Fig. 1D), the relationship between RT and true and recalled offer value (Fig. 1E), the proportion of time spent fixating relevant items (Fig. 2B), and the interaction between choice and valence on the proportion fixation time (Fig. 2C). The results of these analyses are reported in each panel as the posterior deviation of every feature’s coefficient from the grand mean of the four feature coefficients, shown with 95% highest-density intervals. Because these four deviations sum to zero by construction, a feature whose HDI exclude zero (vertical dotted line) is reliably different from the average. Note that per-feature estimates are supported by roughly one quarter of the trials that drive the corresponding pooled effect in the main text. Across all 32 feature-by-analysis comparisons, only two 95% HDIs excluded zero: environment was above the grand mean for the relationship between RT and the number of recalled memories (Δ*β* = 0.065, 95% HDI [0.002, 0.127]), and animacy was above the grand mean for the slope of choice on recalled offer value (Δ*β* = 0.329, 95% HDI [0.004, 0.684]). All other deviations were consistent with no feature-specific effects. Overall, these results indicate that the patterns reported in the main text are broadly robust across the four feature dimensions.

**Table S1:** Behavioral regression coefficients. Choice models are logistic (coefficients on the logit scale); RT models are Gaussian with log(RT) as the dependent variable. All predictors were *z*-scored. RE denotes random intercepts and random slopes per participant for all fixed-effect predictors. Bold rows indicate 95% HDIs that exclude zero.

| Term | $\beta$ | SE | 95% HDI |
| --- | --- | --- | --- |
| *Choice ∼ true value + RE* |  |  |  |
| **Intercept** | **0.183** | **0.053** | **[0.077, 0.287]** |
| **True value** | **0.997** | **0.133** | **[0.750, 1.283]** |
| *Choice ∼ recalled value + RE* |  |  |  |
| **Intercept** | **0.205** | **0.062** | **[0.085, 0.326]** |
| **Recalled value** | **1.292** | **0.134** | **[1.053, 1.585]** |
| *$\log(\text{RT}) \sim$ recalled value + recalled value<sup>2</sup> + RE* |  |  |  |
| **Intercept** | **1.706** | **0.127** | **[1.463, 1.964]** |
| Recalled value | 0.010 | 0.033 | [-0.057, 0.075] |
| **Recalled value<sup>2</sup>** | **-0.080** | **0.019** | **[-0.117, -0.043]** |
| *$\log(\text{RT}) \sim$ true value + true value<sup>2</sup> + RE* |  |  |  |
| **Intercept** | **1.617** | **0.118** | **[1.391, 1.868]** |
| True value | -0.005 | 0.022 | [-0.047, 0.038] |
| True value<sup>2</sup> | 0.010 | 0.012 | [-0.014, 0.033] |

**Table S2:** Gaze allocation during decision making. Coefficients from decision × valence models for offer-relevant and offer-irrelevant episode locations. Models were fit to trial-level fixation proportions (either delta from chance or the raw proportion). RE denotes random intercepts and random slopes per participant for all fixed-effect predictors. Bold rows indicate 95% HDIs that exclude zero.

| Term | $\beta$ | SE | 95% HDI |
| --- | --- | --- | --- |
| *Relevant items: $\delta_{chance} \sim dec \times val + RE$* |  |  |  |
| **Intercept** | **+0.051** | **0.016** | **[0.019, 0.080]** |
| **Decision** | **-0.046** | **0.018** | **[-0.080, -0.007]** |
| Valence | -0.032 | 0.022 | [-0.074, 0.013] |
| **Dec. × Val.** | **+0.099** | **0.030** | **[0.039, 0.156]** |
| *Relevant items: $prop \sim dec \times val + RE$* |  |  |  |
| **Intercept** | **+0.313** | **0.017** | **[0.281, 0.345]** |
| **Decision** | **-0.086** | **0.020** | **[-0.125, -0.046]** |
| **Valence** | **-0.056** | **0.024** | **[-0.103, -0.008]** |
| **Dec. × Val.** | **+0.176** | **0.033** | **[0.110, 0.241]** |
| *Irrelevant items: $\delta_{chance} \sim dec \times val + RE$* |  |  |  |
| **Intercept** | **-0.029** | **0.011** | **[-0.051, -0.007]** |
| Decision | -0.009 | 0.013 | [-0.033, 0.018] |
| Valence | -0.010 | 0.020 | [-0.049, 0.028] |
| Dec. × Val. | +0.011 | 0.019 | [-0.027, 0.048] |
| *Irrelevant items: $prop \sim dec \times val + RE$* |  |  |  |
| **Intercept** | **+0.196** | **0.010** | **[0.175, 0.216]** |
| Decision | +0.025 | 0.013 | [-0.001, 0.051] |
| Valence | +0.039 | 0.020 | [-0.001, 0.079] |
| **Dec. × Val.** | **-0.056** | **0.020** | **[-0.095, -0.017]** |

**Table S3:** Comparison of human participants versus each RNN variant. For each measure, the network benchmark 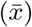 is the mean across 5 simulation seeds. The posterior mean difference (*δ* = Human −RNN), its SE, and its 95% highest density interval (HDI) are reported. Bold rows indicate 95% HDIs that exclude zero.

| Measure | $\bar{x}$ | $\delta$ | SE | 95% HDI |
| --- | --- | --- | --- | --- |
| **Prior memory network** |  |  |  |  |
| Choice accuracy (%) | 0.643 | +0.029 | 0.018 | [−0.005, 0.065] |
| Proportion relevant fixation time | 0.585 | −0.014 | 0.013 | [−0.040, 0.011] |
| Decision × valence prop. relevant | 0.219 | −0.039 | 0.038 | [−0.112, 0.038] |
| Initial fixation prop. relevant | 0.552 | +0.013 | 0.012 | [−0.011, 0.036] |
| Revisit fixation prop. relevant | 0.576 | −0.007 | 0.026 | [−0.061, 0.043] |
| AUC (cumulative prop. fixation time) | 0.937 | −0.023 | 0.013 | [−0.049, 0.002] |
| Cumulative 80% (cumulative prop. fixation time) | 5.800 | +1.392 | 0.867 | [−0.283, 3.056] |
| Total fixations per trial | 6.236 | +2.216 | 1.125 | [−0.021, 4.387] |
| Fixation count × relevance | — | +0.302 | 0.223 | [−0.144, 0.736] |
| **No prior memory network** |  |  |  |  |
| **Choice accuracy (%)** | **0.634** | **+0.037** | **0.018** | **[0.003, 0.071]** |
| Proportion relevant fixation time | 0.563 | +0.008 | 0.013 | [−0.018, 0.037] |
| **Decision × valence prop. relevant** | **0.289** | **−0.108** | **0.038** | **[−0.180, −0.033]** |
| **Initial fixation prop. relevant** | **0.526** | **+0.039** | **0.012** | **[0.015, 0.062]** |
| Revisit fixation prop. relevant | 0.552 | +0.016 | 0.025 | [−0.034, 0.065] |
| AUC (cumulative prop. fixation time) | 0.900 | +0.014 | 0.013 | [−0.012, 0.039] |
| Cumulative 80% (cumulative prop. fixation time) | 8.400 | −1.198 | 0.867 | [−2.983, 0.483] |
| Total fixations per trial | 9.014 | −0.419 | 0.967 | [−2.272, 1.506] |
| **Fixation count × relevance** | — | **+0.442** | **0.220** | **[0.014, 0.874]** |

**Table S4:** Position-based gaze effects for human participants. Top: log fixation duration as a function of relevance, fixation position, and their interaction. Middle: proportion of fixations to relevant locations (minus 0.5) as a function of position. Bottom: proportion of fixations to positive vs. negative reward locaitons as a function of decision type, valence, and position. RE denotes random intercepts and random slopes per participant for all fixed-effect predictors. Bold rows indicate 95% HDIs that exclude zero.

| Term | $\beta$ | SE | 95% HDI |
| --- | --- | --- | --- |
| $\log(dur) \sim relevance \times position + RE$ | | | |
| **Intercept** | **5.810** | **0.061** | **[5.691, 5.933]** |
| **Relevance** | **+0.170** | **0.045** | **[0.080, 0.258]** |
| Position | +0.002 | 0.007 | [-0.012, 0.015] |
| Relevance × Position | -0.012 | 0.008 | [-0.027, 0.004] |
| $\delta_{chance} \sim position + RE$ | | | |
| **Intercept** | **+0.063** | **0.017** | **[0.029, 0.096]** |
| Position | -0.005 | 0.004 | [-0.013, 0.004] |
| $\delta_{chance} \sim dec \times val \times pos + RE$ | | | |
| **Intercept** | **+0.130** | **0.009** | **[0.112, 0.148]** |
| Decision | +0.003 | 0.018 | [-0.034, 0.039] |
| Valence | -0.017 | 0.019 | [-0.056, 0.020] |
| **Position** | **-0.010** | **0.002** | **[-0.015, -0.006]** |
| **Decision × Valence** | **+0.115** | **0.036** | **[0.047, 0.185]** |
| Decision × Position | -0.001 | 0.004 | [-0.009, 0.007] |
| **Valence × Position** | **+0.011** | **0.004** | **[0.003, 0.019]** |
| **Dec. × Val. × Pos.** | **-0.017** | **0.008** | **[-0.033, -0.001]** |

**Table S5:** Fixation-position models: human participants versus each RNN variant. Top: proportion of fixations to relevant locations (minus 0.5) as a function of position. Bottom: proportion of fixations to positive vs. negative reward locations as a function of decision type, valence, and position. RE denotes random intercepts and random slopes per participant for all fixed-effect predictors. Bold rows indicate 95% HDIs that exclude zero.

| Term | $\beta$ | SE | 95% HDI |
| --- | --- | --- | --- |
| **Prior memory network** |  |  |  |
| *Prop. relevant: $\delta \sim position + RE$* |  |  |  |
| Intercept | -0.013 | 0.018 | [-0.048, 0.023] |
| Position | +0.001 | 0.005 | [-0.009, 0.010] |
| *Valence: $\delta \sim 1 + dec \times val \times pos + RE$* |  |  |  |
| Intercept | -0.008 | 0.010 | [-0.028, 0.012] |
| Decision | +0.006 | 0.020 | [-0.032, 0.045] |
| Valence | -0.008 | 0.024 | [-0.055, 0.038] |
| Position | +0.001 | 0.002 | [-0.004, 0.006] |
| Decision × Valence | +0.035 | 0.038 | [-0.039, 0.110] |
| Decision × Position | <0.001 | 0.004 | [-0.008, 0.009] |
| **Valence × Position** | **+0.012** | **0.004** | **[0.004, 0.021]** |
| Dec. × Val. × Pos. | -0.014 | 0.009 | [-0.031, 0.003] |
| **No prior memory network** |  |  |  |
| *Prop. relevant: $\delta \sim position + RE$* |  |  |  |
| **Intercept** | **+0.069** | **0.018** | **[0.034, 0.104]** |
| **Position** | **-0.010** | **0.005** | **[-0.020, -0.001]** |
| *Valence: $\delta \sim 1 + dec \times val \times pos + RE$* |  |  |  |
| **Intercept** | **+0.033** | **0.010** | **[0.014, 0.052]** |
| Decision | +0.006 | 0.019 | [-0.031, 0.045] |
| Valence | -0.017 | 0.024 | [-0.063, 0.031] |
| Position | -0.004 | 0.002 | [-0.009, <0.001] |
| Decision × Valence | +0.003 | 0.038 | [-0.072, 0.079] |
| Decision × Position | <0.001 | 0.004 | [-0.009, 0.008] |
| **Valence × Position** | **+0.013** | **0.004** | **[0.004, 0.021]** |
| Dec. × Val. × Pos. | -0.011 | 0.009 | [-0.028, 0.006] |

**Table S6:** Standalone fixation-position models for RNN variants. Top: log fixation duration as a function of relevance and position. Middle: proportion of fixations to relevant locations (minus 0.5) as a function of position. Bottom: proportion of fixations to positive vs. negative reward locations as a function of decision type, valence, and position. Bold rows indicate 95% HDIs that exclude zero.

| Term | $\beta$ | SE | 95% HDI |
| --- | --- | --- | --- |
| **Prior memory network** |  |  |  |
| $\log(dur) \sim rel \times pos$ | | | |
| **Intercept** | **+0.113** | **0.001** | **[0.111, 0.114]** |
| **Relevance** | **+0.031** | **0.001** | **[0.029, 0.033]** |
| Position | <0.001 | <0.001 | [<0.001, 0.001] |
| **Relevance × Position** | **-0.001** | **<0.001** | **[-0.002, -0.001]** |
| $\delta_{chance} \sim 1 + pos$ | | | |
| **Intercept** | **+0.074** | **0.003** | **[0.069, 0.079]** |
| **Position** | **-0.005** | **0.001** | **[-0.006, -0.004]** |
| $\delta \sim dec \times val \times pos$ | | | |
| **Intercept** | **+0.116** | **0.002** | **[0.113, 0.119]** |
| Decision | -0.004 | 0.003 | [-0.010, 0.002] |
| Valence | -0.002 | 0.003 | [-0.009, 0.004] |
| **Position** | **-0.011** | **<0.001** | **[-0.012, -0.010]** |
| **Decision × Valence** | **+0.059** | **0.006** | **[0.046, 0.072]** |
| Decision × Position | <0.001 | 0.001 | [-0.001, 0.002] |
| Valence × Position | -0.001 | 0.001 | [-0.002, 0.001] |
| **Dec. × Val. × Pos.** | **+0.004** | **0.001** | **[0.001, 0.007]** |
| **No prior memory network** |  |  |  |
| $\log(dur) \sim rel \times pos$ | | | |
| **Intercept** | **+0.120** | **0.001** | **[0.119, 0.122]** |
| **Relevance** | **+0.016** | **0.001** | **[0.015, 0.018]** |
| **Position** | **-0.001** | **<0.001** | **[-0.002, -0.001]** |
| **Relevance × Position** | **+0.002** | **<0.001** | **[0.002, 0.003]** |
| $\delta_{chance} \sim 1 + pos$ | | | |
| **Intercept** | **-0.012** | **0.002** | **[-0.015, -0.009]** |
| **Position** | **+0.008** | **<0.001** | **[0.007, 0.008]** |
| $\delta \sim dec \times val \times pos$ | | | |
| **Intercept** | **+0.042** | **0.002** | **[0.039, 0.046]** |
| Decision | -0.005 | 0.004 | [-0.013, 0.002] |
| Valence | +0.005 | 0.003 | [-0.002, 0.012] |
| Position | <0.001 | <0.001 | [-0.001, 0.001] |
| **Decision × Valence** | **+0.048** | **0.007** | **[0.035, 0.062]** |
| Decision × Position | <0.001 | 0.001 | [-0.001, 0.002] |
| Valence × Position | -0.001 | 0.001 | [-0.002, 0.001] |
| **Dec. × Val. × Pos.** | **+0.008** | **0.002** | **[0.005, 0.011]** |

**Table S7:** Next-fixation location prediction conditional logistic regression coefficients. Fixed-effect posterior means and 95% HDIs for the ten *z*-scored predictors of a conditional logistic regression model (**Figs. 5** and **S5**), fit to human participants, the prior-memory network, and two null models. Predictors are grouped into the three categories shown in **Fig. 5**. Each row reports the posterior mean and 95% HDI. Rows whose HDI excludes zero are shown in bold.

| Predictor | Fit | $\beta$ | 95% HDI |
| --- | --- | --- | --- |
| *Resource-rational* |  |  |  |
| Inverse relevant fixation time | Human | **+0.061** | **[0.033, 0.089]** |
|  | RNN | **+0.115** | **[0.112, 0.118]** |
|  | Adjacent | +0.004 | [−0.003, 0.011] |
|  | Random | 0.000 | [−0.006, 0.005] |
| Clockwise distance | Human | **−0.069** | **[−0.101, −0.039]** |
|  | RNN | **−0.045** | **[−0.048, −0.042]** |
|  | Adjacent | **−1.981** | **[−2.001, −1.962]** |
|  | Random | +0.001 | [−0.005, 0.006] |
| Counter-clockwise distance | Human | **−0.108** | **[−0.151, −0.066]** |
|  | RNN | **−0.040** | **[−0.043, −0.037]** |
|  | Adjacent | **−1.975** | **[−1.995, −1.955]** |
|  | Random | 0.000 | [−0.006, 0.005] |
| *Encoding order* |  |  |  |
| Forward temporal distance | Human | **−0.077** | **[−0.140, −0.015]** |
|  | RNN | +0.001 | [−0.005, 0.007] |
|  | Adjacent | +0.001 | [−0.012, 0.016] |
|  | Random | +0.003 | [−0.010, 0.015] |
| Backward temporal distance | Human | −0.012 | [−0.072, 0.067] |
|  | RNN | −0.001 | [−0.008, 0.005] |
|  | Adjacent | −0.004 | [−0.018, 0.011] |
|  | Random | −0.003 | [−0.015, 0.008] |
| Primacy | Human | −0.001 | [−0.049, 0.042] |
|  | RNN | −0.001 | [−0.005, 0.003] |
|  | Adjacent | +0.002 | [−0.008, 0.012] |
|  | Random | +0.001 | [−0.007, 0.010] |
| Recency | Human | +0.021 | [−0.022, 0.064] |
|  | RNN | +0.002 | [−0.002, 0.007] |
|  | Adjacent | +0.002 | [−0.007, 0.012] |
|  | Random | −0.004 | [−0.012, 0.004] |
| *Biases* |  |  |  |
| Reward magnitude | Human | +0.019 | [−0.018, 0.054] |
|  | RNN | **−0.019** | **[−0.022, −0.016]** |
|  | Adjacent | −0.002 | [−0.009, 0.004] |
|  | Random | +0.002 | [−0.004, 0.008] |
| Signed reward | Human | +0.015 | [−0.018, 0.044] |
|  | RNN | 0.000 | [−0.003, 0.002] |
|  | Adjacent | +0.002 | [−0.004, 0.009] |
|  | Random | −0.001 | [−0.006, 0.005] |
| Previous item | Human | **+0.088** | **[0.027, 0.144]** |
|  | RNN | **−0.127** | **[−0.130, −0.125]** |
|  | Adjacent | **+0.075** | **[0.070, 0.080]** |
|  | Random | −0.002 | [−0.008, 0.003] |

**Table S8:**
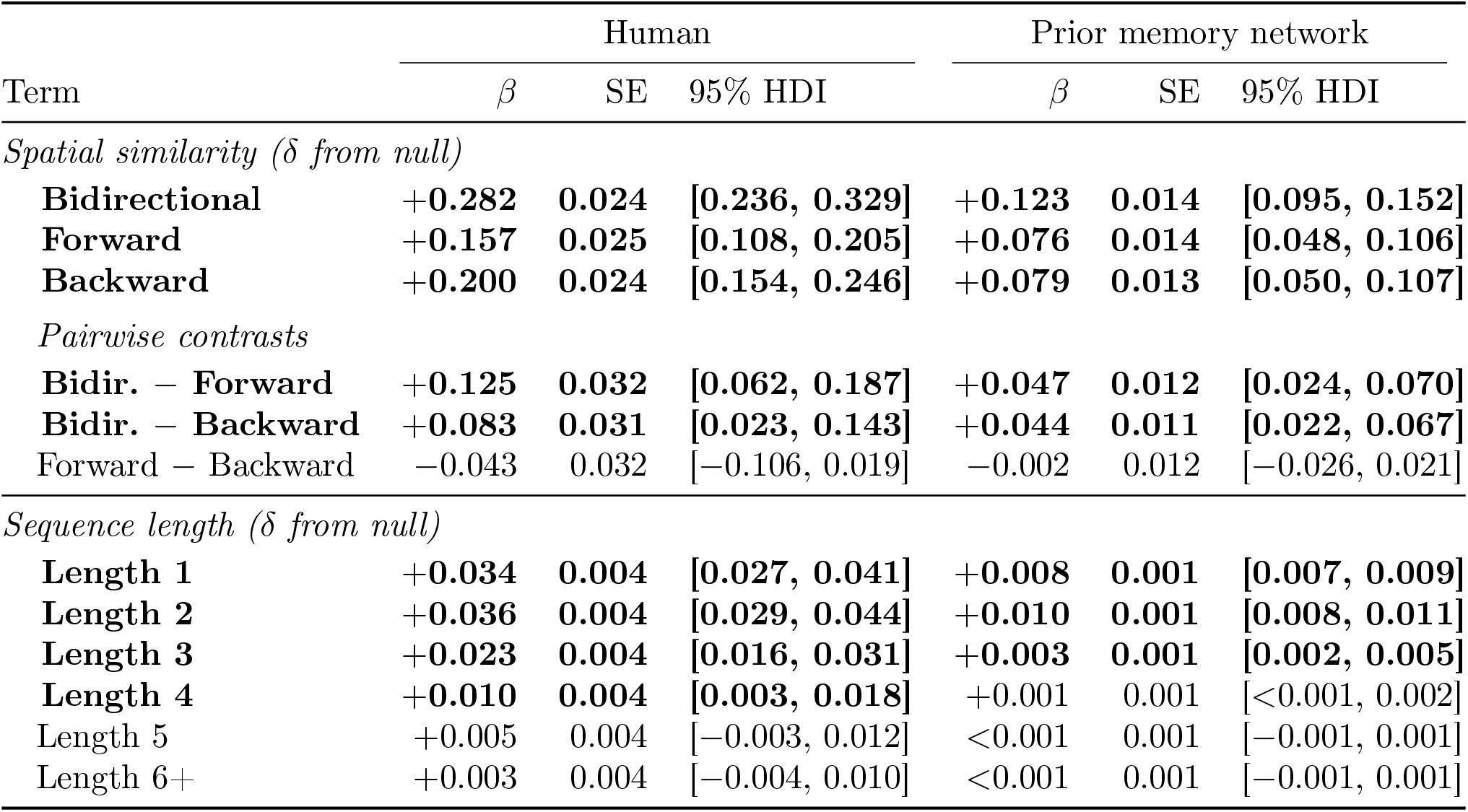
Effects of space on episodic sampling for human participants and the *prior memory* network. Top: spatial similarity to directional sweep templates (*δ* from shuffled null), including pairwise contrasts. Bottom: proportion of transitions in consecutive directional sequences of each length (*δ* from shuffled null). Bold rows indicate 95% HDIs that exclude zero.

**Table S9:** No effects of round on behavioral and eyetracking results. For each primary effect reported in the main text, we refit the corresponding model with round number (as a centered continuous predictor) interacting with the effect of interest as both a fixed effect and a random slope. Each row reports the posterior mean, standard error, and 95% HDI of the interaction coefficient. RE denotes random intercepts and random slopes per participant for every fixed-effect predictor. No interaction was reliably different from zero, indicating that each reported effect was stable across the seven rounds of the experiment.

| Model | Term | $\beta$ | SE | 95% HDI |
| --- | --- | --- | --- | --- |
| Correct $\sim 1 + \text{round} + \text{RE}$ | round | +0.017 | 0.027 | [−0.037, 0.071] |
| $\log(\text{RT}) \sim \text{memories} \times \text{round} + \text{RE}$ | memories × round | +0.007 | 0.013 | [−0.020, 0.032] |
| Choice $\sim \text{recalled value} \times \text{round} + \text{RE}$ | recalled value × round | +0.026 | 0.062 | [−0.094, 0.152] |
| $\log(\text{RT}) \sim (\text{rec. val.} + \text{rec. val.}^2) \times \text{round} + \text{RE}$ | rec. val.<sup>2</sup> × round | +0.005 | 0.008 | [−0.011, 0.021] |
| Relevant fix. prop. $\sim 1 + \text{round} + \text{RE}$ | round | −0.001 | 0.004 | [−0.008, 0.007] |
| Fix. prop. $\sim \text{dec} \times \text{val} \times \text{round} + \text{RE}$ | dec × val × round | +0.013 | 0.011 | [−0.010, 0.035] |
| Item recall prop. $\sim 1 + \text{round} + \text{RE}$ | round | +0.005 | 0.005 | [−0.006, 0.015] |
| True value $\sim \text{recalled value} \times \text{round} + \text{RE}$ | recalled value × round | −0.009 | 0.010 | [−0.029, 0.011] |
| Location recall $\sim 1 + \text{round} + \text{RE}$ | round | +0.202 | 0.209 | [−0.137, 0.672] |

